# Heterogeneous local structures of the microtubule lattice revealed by cryo-ET and non-averaging analysis

**DOI:** 10.1101/2024.04.30.591984

**Authors:** Hanjin Liu, Hiroshi Yamaguchi, Masahide Kikkawa, Tomohiro Shima

**Affiliations:** Department of Biological Sciences, Graduate School of Science, The University of Tokyo Hongo 7-3-1, Bunkyo-ku, Tokyo #113-0033, JAPAN; Department of Cell Biology and Anatomy, Graduate School of Medicine, The University of Tokyo Hongo 7-3-1, Bunkyo-ku, Tokyo #113-0033, JAPAN

**Keywords:** Cytoskeleton, Polymorphism, Heterogeneity, Dynamic instability, GTP-cap

## Abstract

Microtubule cytoskeletons play pivotal roles in various cellular processes, including cell division and locomotion, by dynamically changing their length and distribution in cells through tubulin polymerization/depolymerization. Recent structural studies have revealed the polymorphic lattice structure of microtubules closely correlate with the microtubule dynamics, but the studies were limited to averaged structures. To reveal the transient and localized structures, such as GTP-cap, we developed several non-averaging methods for cryogenic electron tomography to precisely measure the longitudinal spacing and helical twisting of individual microtubule lattices at the tubulin subunit level. Our analysis revealed that polymerizing and depolymerizing ends share a similar structure with regards to lattice spacing. The most distinctive property specific to the polymerizing plus end was left-handed twisting in the inter-dimer interface, suggesting that the twisting might accelerate fast polymerization at the plus ends. Our analysis uncovered the heterogeneity of native microtubules and will be indispensable for the study of microtubules dynamics under physiological contexts or during specific cellular events.

## Introduction

Microtubules are a major cytoskeleton of eukaryotic cells, providing mechanical strength to help cells form their shapes (Gudimchuk and McIntosh, 2021). Microtubules dynamically change their distribution, serving as a platform for motor proteins and microtubule-associated proteins (MAPs) to drive a variety of cellular phenomena including chromosome segregation and cell migration (Kline-Smith and Walczak, 2004). At the molecular level, microtubules are composed of αβ-tubulin dimers assembled into a hollow cylinder shape (Zhang *et al*., 2018), where polymerization and depolymerization are described by the addition and removal of tubulin dimers at both ends (Farmer and Zanic, 2023). Additionally, microtubules stochastically switch between slow polymerization and rapid depolymerization phases (Horio and Hotani, 1986; Cassimeris et al., 1988): otherwise known as dynamic instability. This switching has been thought to be regulated by the nucleotide state at the microtubule tips (Mitchson and Kirschner, 1984): GTP-bound tubulins form stable microtubule lattices immediately after their incorporation into microtubules; the lattice formation triggers GTP hydrolysis, leading to destabilization and depolymerization of the microtubule. As a result, most tubulins in the middle region of a microtubule are thought to be in the unstable GDP-bound state, while freshly incorporated tubulin molecules at the microtubule tips hold GTP to form stable GTP caps. The detection of GTP-tubulins using conformation-specific antibodies further confirmed that the tip regions are in a distinct state (Dimitrov et al., 2008). Regulation of the GTP cap structure, the GTPase activity, and the synergy of MAPs and the microtubule lattice was thought to play a central role in controlling cell shapes and cellular components during cell division and locomotion (Zanic et al., 2013). However, growing interest in the middle regions of microtubules has led to a novel concept of lattice plasticity (Cross, 2019), a concept backed up by several studies showing GTP-tubulin incorporation from the lattice (Aumeier et al., 2016; Vemu et al., 2018) or conformational changes in the middle regions induced by MAPs after polymerization (Shima et al., 2018; Siahaan et al., 2022). These features imply that microtubules should have highly heterogenic lattice structures, which enables them to fulfill diverse cellular functions.

To gain insight into the conformational changes of microtubules, cryogenic electron microscopy (cryo-EM) has revealed various lattice structures of microtubules depending on the microtubule status (Alushin et al., 2014; Zhang et al., 2015; Morikawa et al., 2015; Zhang et al., 2017; Shima et al., 2018; Estévez-Gallego et al., 2020; LaFrance et al., 2022). Interestingly, the lattice structures have shown a strong correlation with the dynamic properties of the microtubules. For instance, depolymerization-resistant microtubules such as microtubules polymerized with the very slowly hydrolysable GTP analogue guanosine-5’-[(α, β)-methyleno]triphosphate (GMPCPP), microtubules treated with anti-cancer drug paclitaxel, or microtubules fully decorated with the motor protein KIF5C take a tubulin lattice with a longitudinally expanded conformation (Alushin et al., 2014, Shima et al., 2018) when compared to the structure of GDP-bound microtubules (GDP-MTs). This observation was confirmed by X-ray diffraction and fluorescence microscopy (Kamimura et al., 2016; Shima et al., 2018; Peet et al., 2018). In vitro reconstitution assays have proven that conversions of the microtubule lattice structure also regulate the binding affinity or catalytical activity of associated proteins (Zanic et al., 2009; Morikawa et al., 2015; Liu and Shima, 2023; Yue et al., 2023). These results strongly suggest that microtubule lattice structures undergo local regulation within cells to appropriately modulate microtubule dynamics and protein binding kinetics (Shima et al., 2018; Siahaan et al., 2022).

Despite advances in structural biology, the characterization of native and/or local microtubule lattice structures, particularly the GTP cap, has been hampered by potential heterogeneity among microtubule subregions. To obtain a high-resolution structure, conventional cryo-EM methods require an adequate number of homogenous GTP caps, which itself is challenging. Even if such images were collected, the length of each GTP cap and the consistency of the structures between individual GTP caps would remain unknown. To overcome this problem, previous studies have resorted to a “structure homogenization” strategy. For a high-resolution analysis, GTP-MT was reconstituted using GMPCPP, a slowly hydrolyzing GTP analog, or the tubulin mutant E254A, which is unable to hydrolyze GTP. The artificial microtubule structures obtained revealed the possibility of lattice expansion in GTP caps (Alushin et al., 2014; LaFrance et al., 2022), and, accordingly, GMPCPP-MT has been extensively used as a proxy for GTP cap in various in vitro reconstitution assays (Guedes-Dias et al., 2019), molecular dynamics (Igaev and Grubmüller, 2022) and energy simulation studies (Manka and Moore, 2018). However, a study using BeF_3_^-^, a strict mimic of the orbital of γ-phosphate, suggested the opposite: GTP caps are not expanded and the lattice expansion in GMPCPP-MT is an artificial effect of the carbon atom located between the α- and β-phosphate of the GMPCPP, not γ-phosphate itself (Estévez-Gallego et al., 2020). The fact that the lattice of yeast microtubules, which undergo dynamic instability as well, is always expanded regardless of the bound nucleotide (Howes et al., 2017) also implies that lattice expansion is not the direct regulator of the microtubule’s dynamic nature. The ongoing controversy underscores that no matter how carefully we mimic the structure, the accuracy of the mimicry can only be confirmed by observing the native structure.

Here, to overcome this problem, we developed novel computational methods for cryogenic electron tomography (cryo-ET) to perform a non-averaged structural analysis of native microtubules. We first used a Fourier transform specialized for cylindric structures to determine lattice helical parameters. This method, named CFT (<u>c</u>ylindric <u>F</u>ourier transformation), can measure the local lattice parameters of individual microtubules. Remarkably, only a 49-nm-long sub-volume (six dimers × all protofilaments) was sufficient to distinguish GMPCPP-MTs from GDP-MTs with >99.9% accuracy. The application of CFT to the microtubule tip regions revealed that polymerizing microtubule tips are not expanded, instead twisting in the left-handed direction. This observation was further confirmed by higher resolution methods to directly estimate the coordinates of each tubulin monomer, which we named Viterbi alignment and restricted mesh annealing (RMA). As these methods are able to measure lattice structure at the near single-molecule level, our analysis identified that lattice twisting at the polymerizing plus ends originates from the inter-tubulin dimer interface and that the mean lattice spacing increases at the curved regions of microtubules. Our novel computational methods unveiled the genuine heterogeneous structures of intact microtubule tips and curved regions, shedding light on the role of such local lattice structures. These methods have the potential to provide new insights into how the dynamic instability and MAP binding are controlled by the local structures of individual microtubules, thereby leading to intricate cell behaviors.

## Results

### Cylindric Fourier transformation accurately estimates microtubule lattice structures with 49-nm fragments

First, we tried to measure the lattice structure in a narrow region of a single microtubule using the inherent periodicity of the microtubule structure. To measure the periodicity, Fourier transformation has already been used with cryo-EM and X-ray diffraction (Estévez-Gallego et al., 2020; Ku et al., 2020; Guyomar et al., 2022). We speculated that we can further leverage the power of Fourier transformation by cylindric coordinate transformation and up-sampling of the power spectra to maximize the precision of the peak detection (see Materials and Methods). We named this workflow CFT (<u>c</u>ylindric <u>F</u>ourier transformation) and referred to the CFT analysis of a short microtubule fragment as local-CFT and that of the entire microtubule as global-CFT. Note that global-CFT requires the straightening of microtubule sub-volumes, but local-CFT does not, owing to the large persistent length of a microtubule (∼300 μm, Wisanpitayakorn et al., 2022). To validate our CFT analysis, we collected and analyzed cryo-ET images of GDP-MTs (Fig. 1A) and GMPCPP-MTs (Fig. 1B), which are known to be different in their lattice spacing (longitudinal distance between tubulin monomers) and the twist angle (right-handed rotation of lattice per monomer). It is also known that the N_S lattice type (N: number of protofilaments; S: number of longitudinal monomer-shifts per helix turn) affects the microtubule lattice parameters; thus, we took the N_S type into account for the later analysis. For precise measurements, the pixel size was calibrated by the local-CFT analysis of tobacco mosaic virus, a cylindrical periodic structure with an ∼2.3-nm helical pitch (Fig. S1). The cylindric power spectrum of microtubules resembles the power spectrum of tubulin sheets (Portran et al., 2017), indicating that CFT successfully extracted the periodic patterns of the lattice. When we looked at the peaks corresponding to the lattice spacing, we noticed that 49-nm windows (six dimers × all protofilaments) is sufficient to separate the peaks of GDP-MT and GMPCPP-MT (Fig. S2). We also measured lattice spacing with different sizes of microtubule fragments, finding the accuracy of detection was >99.9% with a 49-nm window (Fig. S3). This result demonstrates that CFT can accurately measure lattice parameters.

**Fig. 1:**
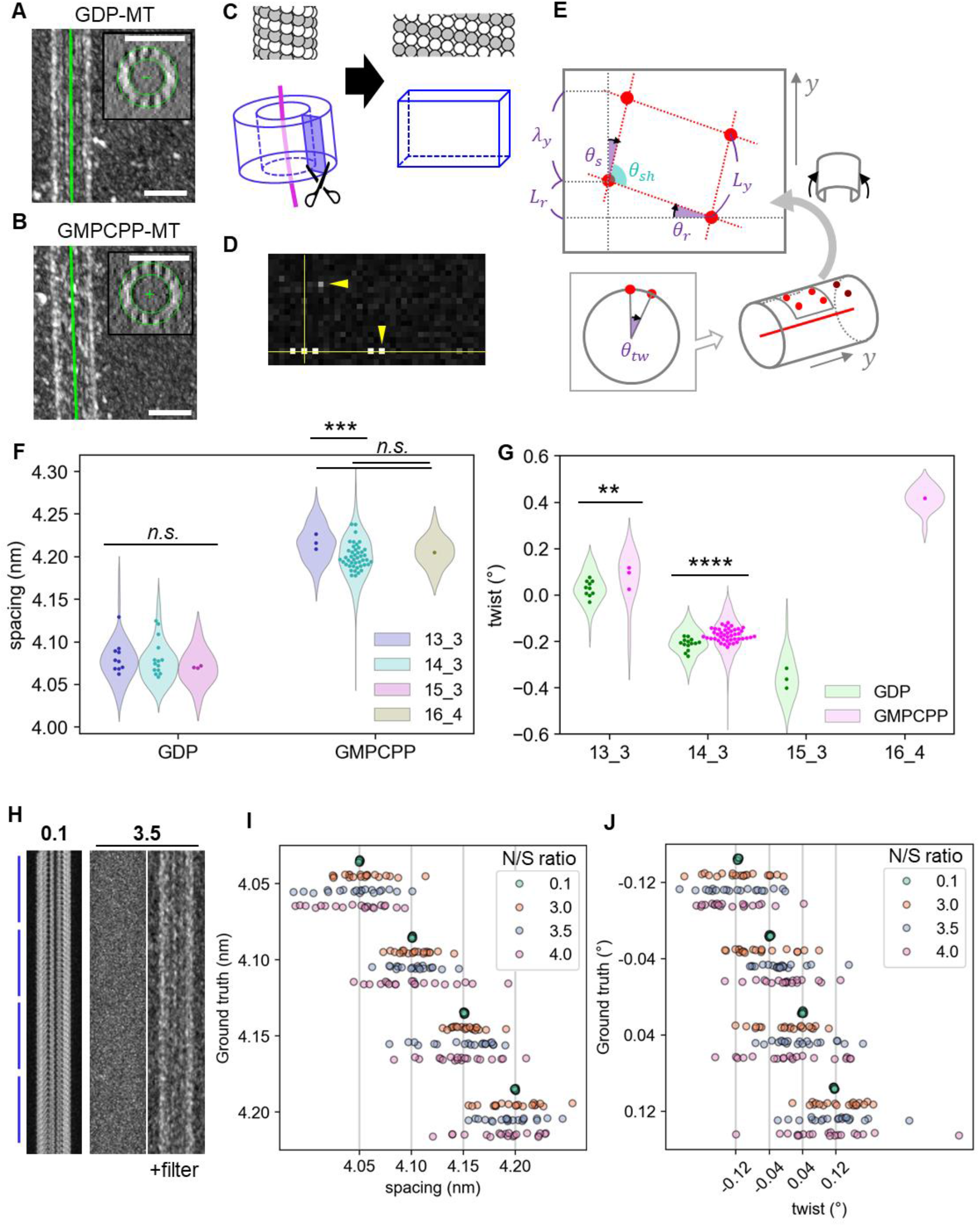
Application of CFT to quantify microtubule lattice parameters. (A, B) Tomograms of GDP-MT and GMPCPP-MT. Tomograms are binned by 4x4x4 pixels and low-pass filtered to make the microtubules visible. Green curves are the fitted spline. Insets are projections of a 49-nm microtubule fragment along the spline. Scale bars, 25 nm. (C) Schematic illustration of the cylindric coordinate transformation along a microtubule. (D) A representative power spectrum calculated by CFT of a 49-nm microtubule fragment. (E) Schematic illustration of the definition of the lattice parameters. *λ*_*y*_: pitch length, *L*_*y*_: spacing, *L*_*r*_: rise length, θ_*r*_: rise angle, θ_*s*_: skew angle, θ_*tw*_ : twist angle, θ_*sh*_ : shear angle. (F, G) Lattice parameter quantification of GDP-MT and GMPCPP-MT focusing on (F) the longitudinal lattice spacing and (G) the twist angle. **: *p* < 0.01, ***: *p* < 0.001, ****: *p* < 0.0001 (Tukey’s HSD test). (H) Simulated tomograms of microtubules with piecewise lattice expansion corresponding to (I) with low-level (N/S = 0.1) and experimental-level (N/S = 3.5) noise. (I, J) CFT analysis of the simulated tomograms. Microtubules with (I) lattice spacing spanning from 4.05 nm to 4.20 nm or (J) twists spanning from -0.12° to 0.12° were used for the simulation. y-axis, true values; x-axis, estimated values.

To verify the accuracy of this method, the lattice parameters of each microtubule were measured. We fitted microtubules to spline curves based on manual drawing and image alignment and then quantified the lattice parameters of 49-nm microtubule fragments using local-CFT (Table S1, Fig. 1A-E). The results showed that CFT recapitulated all known features. The mean spacings were 4.21 nm for GMPCPP-MT and 4.08 nm for GDP-MT independent of the protofilament number (Fig. 1F, Table S1), consistent with previous structural analyses that concluded the lattice spacing of GMPCPP-MT is ∼3% longer than that of GDP-MT (Alushin et al., 2014; Kamimura et al., 2016). The mean twist was 0.03° for 13_3 GDP-MT and -0.20° for 14_3 GDP-MT (Fig. 1G); these values are again consistent with previous studies (Zhang et al., 2015; Zhang et al., 2018; LaFrance et al., 2022; Zhou et al., 2023). 15_3 GDP-MTs showed even more strong twists (−0.36°; Table S1, Fig. 1G), as reported in previous negative staining work (Chrétien and Fuller, 2000). Note that previous studies usually calculated the dimer twist, which is twice the monomer twist examined in this study. Although the conventional twist angle calculation based on Moiré patterns is not suitable for weakly twisted 13_3 GDP-MTs (Estévez-Gallego et al., 2020), local-CFT is equally applicable to all types of microtubules. CFT analysis can also measure the skew angle (another representation of twisting), rise angle (angle of the adjacent tubulin in the neighbor protofilaments) and the rise length (rise angle represented as a distance) (see Fig. 1E for the definition). The measured rise length was 0.942 nm for 13_3 GDP-MT and 0.871 nm for 14_3 GDP-MT, which are consistent with a previous study (Zhang et al., 2015, in which they called the rise length a “3-start rise”). Using these parameters, we calculated the start number (the S of N_S microtubules), which is 3 with few exceptions. Thus, the analysis shows the accuracy of CFT in determining local lattice parameters.

To further check the robustness of CFT, we simulated tomograms of microtubules by computational projections of tubulins, noise addition and back-projections (Fig. S4A, Fig. 1H). With the experimentally acquired images, the noise level to be added to the simulated tilt series was estimated to be 3.5× the maximum intensity of the tubulin signal (Fig. S4B, C). Using 13_3 microtubules with piecewise expansion or twisting, the estimation error of 49-nm CFT was ∼0.023 nm for the lattice spacing and ∼0.064° for the twist angle in terms of the standard deviation (*σ*) from the corresponding ground truths (Table S2, S3; Fig. 1I, 1J). CFT is also applicable to 14_3 microtubules, although they adopt twist angles in a distinct range from 13_3 microtubules (Table S4, Fig. S4D). We also checked if CFT returns a biased estimation with different microtubule orientations and curvatures, assuming that tubulins experience expansion and compaction equally at the curved regions (Fig. S4E, F). These simulations confirmed that CFT correctly estimates lattice parameters as far as a microtubule within 60° relative to the tilt series rotation axis and local curvature <1.5 μm^-1^. Results of the CFT analysis on the experimental and simulated microtubules strongly support that CFT is appropriate for the quantification of individual microtubule local structures, including spacing, twist angle and rise angle.

### Microtubule polymerizing tips are not stabilized by lattice expansion

The results shown above indicate that CFT can be applied to study transient local structures of dynamic microtubules such as GTP cap. To get the genuine structural profiles of dynamic microtubule ends, we prepared cryo-ET grids with polymerizing and depolymerizing microtubules (Fig. 2A). Instead of polymerizing with GMPCPP seed microtubules, we spontaneously nucleated microtubules from soluble tubulins to make native microtubules. Tomograms showed that the structures of the microtubule lattice were highly diverse, as previously reported (Atherton et al., 2018). Although the majority was 13_3 and 14_3 microtubules, we also found 12_3, 14_2, 15_3 and 16_3 microtubules (Fig. S5A). The ratio of 13_3 microtubules to 14_3 microtubules was much smaller in the depolymerizing samples (Fig. S5A), probably because 13_3 microtubules undergo catastrophe more frequently (Rai et al., 2021), which makes them less stable at the onset of tubulin washout. Notably, there were a few 14_3 microtubules with positive twists (referred to as 14*_3 microtubules hereafter) (Fig. S5B), and one of them steeply turned to the standard negatively-twisted lattice conformation, suggesting that there is no lattice state allowed between 14_3 and 14*_3 microtubules and that the 14*_3 structure is metastable. Therefore, we excluded 14*_3 microtubules from the following analysis and focused on 13_3 and 14_3 microtubules. We also found the 13-to-14 protofilament number transition (Fig. S5C) and that the lattice parameter changed in response to the transition (Fig. S5D). At the transition point, the change in the rise angle was one step, but the twist angle dropped from ∼0° to ∼-0.6° before changing to the ordinary value ∼-0.3° for the 14_3 lattice, suggesting that the transition points take a unique structure rather than a simple mixture of two structures.

**Fig. 2:**
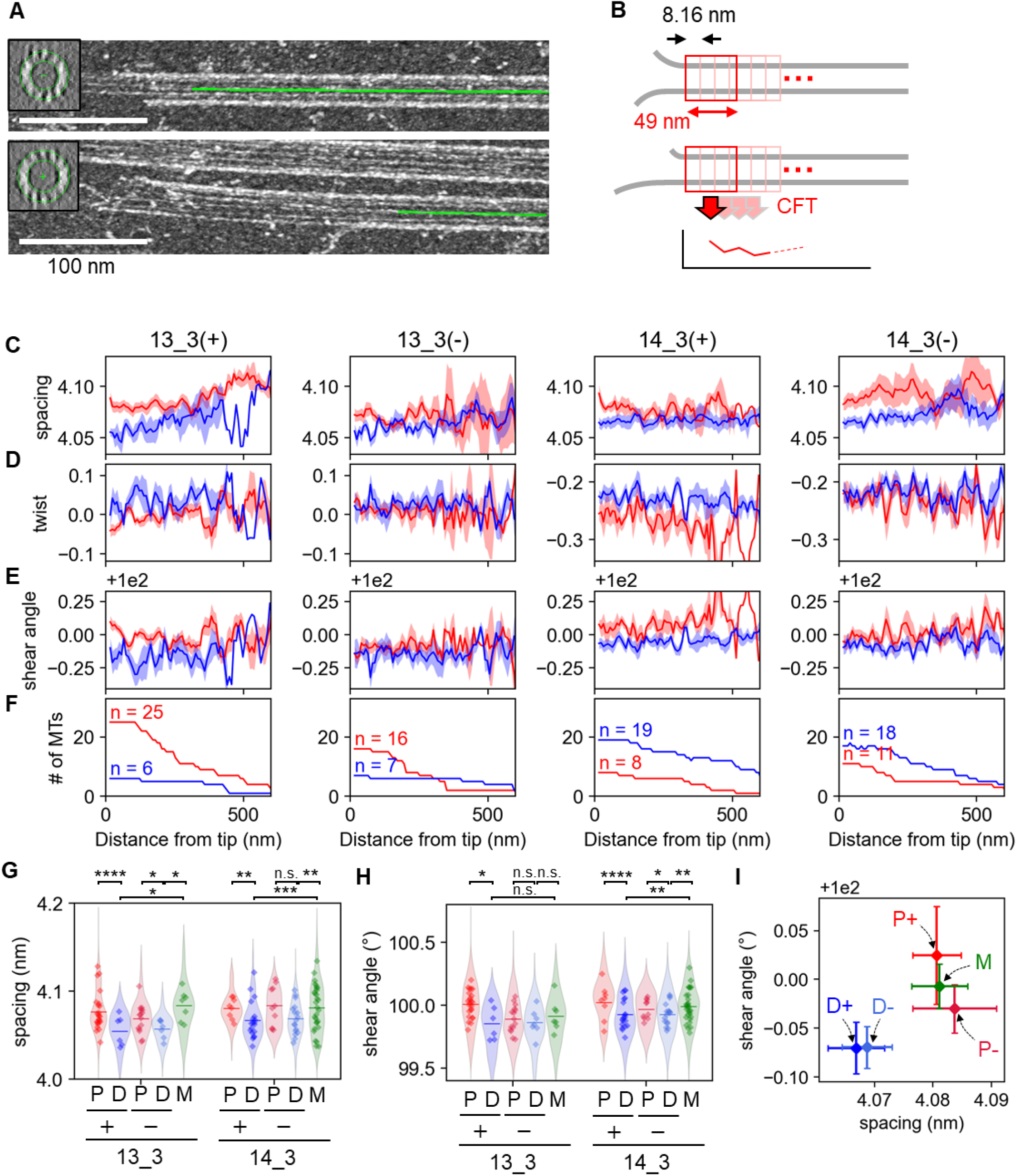
CFT analysis of polymerizing and depolymerizing microtubule tips. (A) Representative microtubule polymerizing (upper) and depolymerizing (lower) tips. Tomograms were binned by 4×4×4 pixels and low-pass filtered to make the microtubules visible. Insets are 49-nm projections of a microtubule fragment along the spline. (B) Overview of the distance-wise aggregation of parameters. (C-F) Profiles of the lattice parameters or sample sizes along microtubules of different types (red, polymerizing microtubule tips; blue, depolymerizing microtubule tips). Solid lines and shaded areas are means ± S.E. Lattice spacing (C), twist angle (D), lattice shear angle (E) and sample size (F). (G, H) Plot of the spacing (G) and shear angle (H) in a 200-nm region from the tips of the microtubules. The dots and violin plots show the mean of each microtubule and the distribution of the values of 49-nm fragments, respectively. P: polymerizing, D: depolymerizing, M: middle region. *: *p* < 0.05, **: *p* < 0.01, ***: *p* < 0.001, ****: *p* < 0.0001 (Welch’s t test). (I) 2D plot of (G) and (H). Data points for 13_3 and 14_3 microtubules were concatenated. Error bars are means ± S.E.

Then, we compared the lattice parameters of the polymerizing and depolymerizing ends to answer the long-standing question of whether the lattice of GTP caps is expanded. To get the consensus profile of the microtubule lattice, we measured 49-nm local lattice parameters along microtubules for every fragment at 8.16-nm intervals, grouped all the profiles by the N_S lattice types, and averaged the parameters for each local region from the tips (Fig. 2B–2F). To avoid microtubule bending from affecting the results, microtubule fragments with a local curvature larger than 0.4 μm^-1^ were excluded from the averages, as mean spacing increased beyond that range (Fig. S6A). Note that every microtubule has a different analyzable length, which caused the sample size to decrease as the distance increased (Fig. 2F). We first focused on the lattice spacing to resolve the controversial conclusions between studies using different methods to mimic GTP caps (GMPCPP in Alushin et al., 2014; GDP-BeF_3_^-^ in Estévez-Gallego et al., 2020; and E254A mutation in LaFrance et al., 2022). If the lattice of GTP caps is expanded, as suggested by Alushin et al. and LaFrance et al., our polymerization condition (48 μM tubulin, 1 mM GTP, 37°C) should produce microtubules with expanded lattices at their tips. Nevertheless, profiles measured by local-CFT did not show any increase in the lattice spacing at the polymerizing tips. Instead, the lattice spacing of polymerizing tips was constantly and slightly larger than the depolymerizing ones (Fig. 2C). To quantitatively compare the lattice spacings of polymerizing and depolymerizing microtubules, we collected all the local-CFT parameters in the range of 0 – 200 nm from the tips (Fig. 2G). As a control, we analyzed the microtubule middle regions that were acquired in the same cryo-ET grids as depolymerizing microtubules, which are supposed to be the ground state of the GDP-MT lattice. Interestingly, the lattice spacings of polymerizing tips and middle regions were longer than those of depolymerizing tips (∼4.08 nm compared to ∼4.06 nm), suggesting that lattice compaction is a characteristic conformational change proceeding a microtubule catastrophe event (Fig. 2G; Table S5). Note that the relative lattice expansion rate of polymerizing tips and middle regions (∼0.5%) is much smaller than that associated with strong stabilization effects such as GMPCPP-MT, taxol-bound GDP-MT and the E254A mutation (∼3%; Kamimura et al., 2016; LaFrance et al., 2022) and even smaller than the expansion triggered by KIF5C-binding (∼1.2%; Shima et al., 2018), which has a milder stabilization effect. Taken together, we concluded that the lattice of native GTP cap is not expanded, but that of depolymerizing tips is compacted on average.

### Polymerizing plus ends have unique shearing of the microtubule lattice

Next, we focused on the lattice twist. The twist angle, *θ*_*tw*_, is defined by the right-handed rotation to fit a monomer to the next one (Fig. 1E). Same as the microtubule middle regions, both polymerizing and depolymerizing 13_3 microtubules had little twist, while most 14_3 microtubules were negatively twisted (Fig. 2D). Twist angles were shifted by ∼0.05° negatively for polymerizing tips compared to depolymerizing tips, and the shift did not converge to the level of depolymerizing microtubules within the analyzed ranges (Fig. 2D). This trend was consistent for both 13_3 and 14_3 microtubules, but the difference was less obvious for minus ends. It is noteworthy that the mean twist of polymerizing 13-protofilaments plus and minus ends was almost zero, which is similar to the value obtained in the previous average structure of GDP•Pi-MT prepared by fast microtubule nucleation using doublecortin (Manka and Moores, 2018). Furthermore, a negative shift in the twist angle is a common conformational change among microtubules that are considered to mimic the GDP•Pi state, including GTPγS-MT, E254N mutation and GDP•AlF_3_-MT, indicating that γ-phosphate tends to twist microtubule lattices toward the (left-handed) direction.

Since the twist angle depends on both the number of protofilaments in the microtubule and coordinate of tubulin inside the lattice, the contribution of each was investigated. The ∼0.05° change in the twist angle shown above is even smaller than the difference between the mean twist angles of 13_3 and 14_3 microtubules (Table S1). Notably, these microtubules show very similar properties such as growth rate, catastrophe frequency and EB3 end-tracking (Rai et al., 2021). At the molecular level, the property of a microtubule should be determined by the relative positioning of the adjacent tubulins. However, the twist angle is defined by the protofilament orientation relative to the microtubule center axis, which is not relevant to the surface geometry. We noticed that the angle between longitudinal and lateral tubulin neighbors can be calculated from the parameters determined by CFT and that the changes in this angle deform the local molecular structure. We named this angle parameter “shear angle”, because this angle defines the shearing of the parallelograms of the tubulin grid, and analyzed the distribution of local shear angles (Fig. 1E). As expected, depolymerizing microtubules showed similar mean shear angles among 12_3, 13_3, 14_3, 15_3 and 16_3 lattices despite the monotonic decrease in the mean twist angles (Fig. S7). However, the shear angles of the polymerizing ends were consistently larger than the depolymerizing ends: on average, shear angles of 13_3 and 14_3 plus ends were larger by 0.16° and 0.13°, respectively (Table S5; Fig. 2E, H). The mean shear angle of the microtubule middle regions was, like the case for the lattice spacing, closer to the values of polymerizing ends (Fig. 2H). Although only the shear angles of 14_3 microtubules showed statistical significance between the ends and the middle regions of depolymerizing microtubules (Fig. 2H), the 2D plot between spacings and shear angles (Fig. 2I) indicated that the middle regions of depolymerizing microtubules belong to the group of polymerizing ends. These results suggest that fast polymerization at the plus ends involves lattice shearing independent of the number of protofilaments in the microtubules.

### Physical constraints improve subtomogram alignment on microtubules and make it feasible for lattice parameter measurements

The CFT analysis revealed that there are no expanded regions at the polymerizing tips, but due to limitations of the CFT, there remains a possibility that the very tip of the ends is expanded. The CFT cannot measure the parameters of tubulins in the protofilaments protruded from the complete cylindrical region of microtubules and the “resolution” of the local-CFT analysis is 49 nm, which contains the signal from ∼80 tubulin dimers. The CFT results may also be slightly affected by some conformational changes that are not relevant to the periodicity, since the Fourier transform is a superposition of all the signals in the original image. To overcome these drawbacks, we next investigated the possibility of quantifying the lattice structure in real space by directly determining the positional and angular coordinates of tubulin. In cryo-ET image analysis, subtomogram alignment is used to precisely estimate molecule positions to obtain average images (Wan and Briggs, 2016). Although the usefulness of averaging subtomograms is well demonstrated for in situ structural analysis (Sun et al., 2022; Erdmann et al., 2021), there is no evidence that coordinate estimation by conventional alignment algorithms is accurate enough to quantify local microtubule lattice parameters (Fig. S8). To address this question, we analyzed simulated microtubules with 6×10 monomers forming 0.1-nm expanded patches (Fig. 3A) and calculated lattice parameters from the tubulin coordinates determined by aligning the tubulin template to the average of the high-resolution α- and β-tubulin densities of GDP-MT determined by the single particle analysis (Fig. 3B–3D; see Materials and Methods for the real-space quantification of the lattice parameters). Although conventional alignment algorithms have been widely used to align protein molecules to acquire high-resolution images, this algorithms was too noise-sensitive for a local structural analysis (Fig. 3D).

**Fig. 3:**
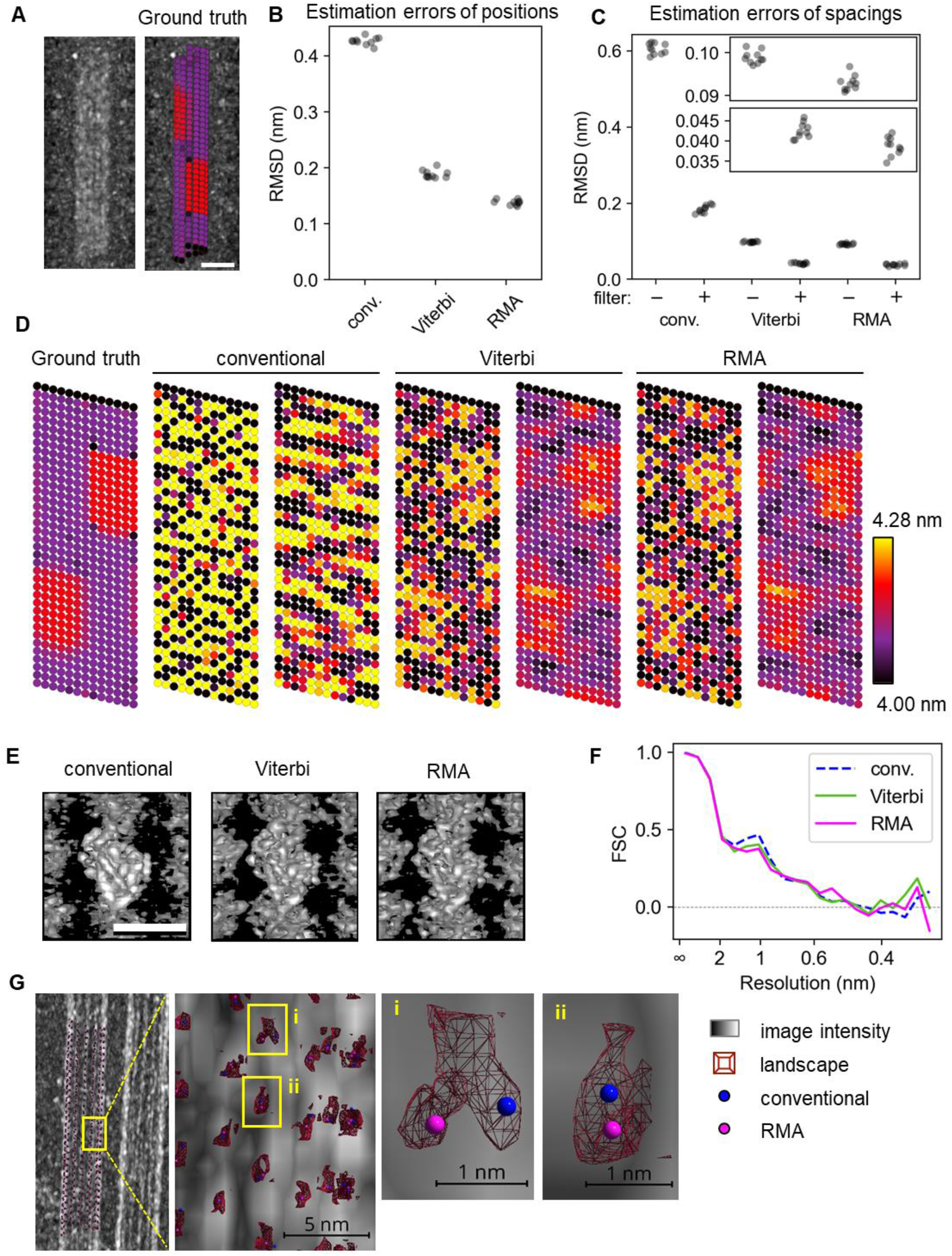
Constrained image alignment algorithms. (A) A representative tomogram of a simulated microtubule, and the 3D view of the ground-truth molecules colored by their lattice spacings. For the color bar, see (D). The tomogram was binned by 4×4×4 pixels and low-pass filtered to make the microtubule visible. (B) Plot of RMSDs of the ground truth and estimated tubulin positions using different alignment methods. (C) Plot of RMSDs of the ground truth and estimated tubulin longitudinal intervals. Insets are zoomed plots. The “filter” indicates a 2×3 mean filter of the interval values. (D) Comparison of the alignment results in flat views from one of the simulations in (B). For each algorithm, the right flat view is the result of mean filtering. Note that the molecules on the top row are painted black because the inter-molecular distance cannot be defined. (E) Subtomogram averages of the tubulins using the results of (D). The threshold levels of the iso-surface are set to the same value. (F) Fourier shell correlation of each alignment. (G) A representative landscape construction on an experimentally acquired tomogram. An arbitrary surface of the landscape is shown as wireframes (the alignment score inside the wireframe is higher than that of the outside). (i) shows a representative optimal coordinate of the conventional alignment being distantly deviated from that of RMA. This deviation may have caused the large RMSD values of the conventional method used in (B) and (C). (ii) shows a representative result of the conventional alignment and RMA being close to each other.

We noticed that this overfitting problem of conventional alignment algorithms may be caused by the high degree of freedom of the tubulin position during alignment. For real microtubules, the displacement of tubulin molecules should be strongly restricted by the physical contact between adjacent tubulins, but in conventional alignment methods, this condition is not considered at all. Therefore, we introduced distance and angle constraints between every set of tubulin neighbors, and the constraints were treated as penalties of the alignment score. However, to get the optimal alignment with constraints, every possible combination of tubulin coordinates has to be used for the score calculation, which results in an exponential increase of computation complexity with the naïve implementation. To solve this optimization problem, we implemented two algorithms: Viterbi alignment, which returns the strictly optimal alignment with longitudinal constraints using the Viterbi algorithm (Viterbi, 1967), and RMA (restricted <u>m</u>esh <u>a</u>nnealing), which approximately solves the optimization problem with both longitudinal and lateral constraints using the simulated annealing algorithm (Fig. S9). Constrained alignment methods up-sample image pixels by five times, resulting in the precision of the tubulin positional coordinates being 0.26 nm (pixel scale) / 5 ≅0.05 nm. To improve the per-molecule resolution, we also implemented a 2×3 (two monomers for each of three protofilaments) mean filter to smooth the calculated lattice structure. Compared to the result of conventional alignment algorithms, Viterbi alignment and RMA showed superior performance in estimating the coordinates and monomer spacings (Fig. 3D–G). With these methods, a 0.1-nm lattice expansion can be distinguished at the near single-molecular level. This improved performance is attributed to the ability of these methods to visualize the expanded patches even without mean filtering. Quantification of the alignment precision using root mean squared deviation (RMSD) showed that constrained alignment algorithms gave much better alignment results than conventional methods, with an RMSD of the spacing after filtering of less than 0.05 nm, which is in stark contrast to the ∼0.2 nm observed with conventional methods (Fig. 3C)). In addition, the results from RMA were slightly better than those from Viterbi alignment (Fig. 3B, C). Accordingly, we decided to use RMA for the following analysis.

### Fresh tubulins have a higher tendency to expand and can remain as expanded clusters

Taking advantage of RMA, we quantified tubulin coordinates in the very tip of both polymerizing/depolymerizing ends, except for the flared regions (Fig. 4A, B). Longitudinal spacings were calculated for every protofilament so that we obtained the mean spacing of the *n*-th monomer from the straightened tip (Fig. 4C). The first tubulin monomers at the polymerizing tip showed slightly longer spacing than the rest of the region. The mean lattice spacing gradually shortened from the tip to the eighth tubulin of each protofilament and remained the same in the subsequent regions. The result suggests that freshly incorporated tubulin tends to form the expanded lattice. However, the expansion rate at the first tubulin from the tip was much lower (∼1.2%) than that expected from GMPCPP-MT (∼3%), suggesting that the majority of tubulin has a compacted lattice even at the very tip of polymerizing microtubules. The results of RMA also showed that the tubulin with an expanded lattice appeared to cluster at the polymerizing ends (Fig. 4B). Therefore, we further investigated whether these local expansions persist after the microtubule has completely formed a cylinder. We ran RMA on the polymerizing and depolymerizing ends, and subtomogram averages using the alignment results indicated that RMA successfully located tubulins at the proper coordinates (Fig. 4D). To quantify the rate of appearance of the expanded clusters, a cluster was defined as a contiguous region of tubulins with lattice spacing above a threshold (so-called morphological image analysis, or the “analyze particle” function in ImageJ). We chose 4.14 nm as the threshold value, which is ∼2.8σ larger than the mean lattice spacing measured by CFT for 13_3 polymerizing plus ends and ∼4σ larger for 14_3 polymerizing plus ends (Table S5). RMA revealed that even depolymerizing ends contain the expanded cluster, indicating the presence of both expanded and compacted lattices in both types of microtubules. The occupancy of the expanded tubulin (the number of tubulins above the threshold divided by the total number) and the frequency of the expanded cluster (appearance per μm) showed a broad distribution in both polymerizing and depolymerizing ends (Fig. 4E), further supporting the notion of a highly heterogeneous microtubule lattice structure, not only in the polymerizing ends but also in the depolymerizing ends. Indeed, the variances of both indicators of depolymerizing microtubules (occupancy: 7.8 ± 7.5%; frequency: 21 ± 19 μm^-1^; mean ± S.D.) were larger than expected from the misalignment due to the experimental noise (Fig. S4; noise-to-signal ratio, N/S = 3.5) and the mean expansion (4.06 nm) of depolymerizing microtubules (Fig. S10; occupancy: 1.1 ± 1.0%; frequency: 8.0 ± 6.0 μm^-1^; mean ± S.D.). The high degree of heterogeneity in the microtubule lattice is also consistent with the low resolution of subtomogram averages before RMA alignment, where constant intermolecular distances are used to initialize molecule coordinates for each microtubule (Fig. 4D). Comparing both ends, the depolymerizing ends contains a lower number of expanded tubulin and clusters, resulting in a lower heterogeneity in the lattice structure. Yet, this structural analysis indicates that microtubules have a heterogenetic lattice consisting of a mosaic-like mixture of the expanded and compacted tubulins in both polymerizing and depolymerizing ends.

**Fig. 4:**
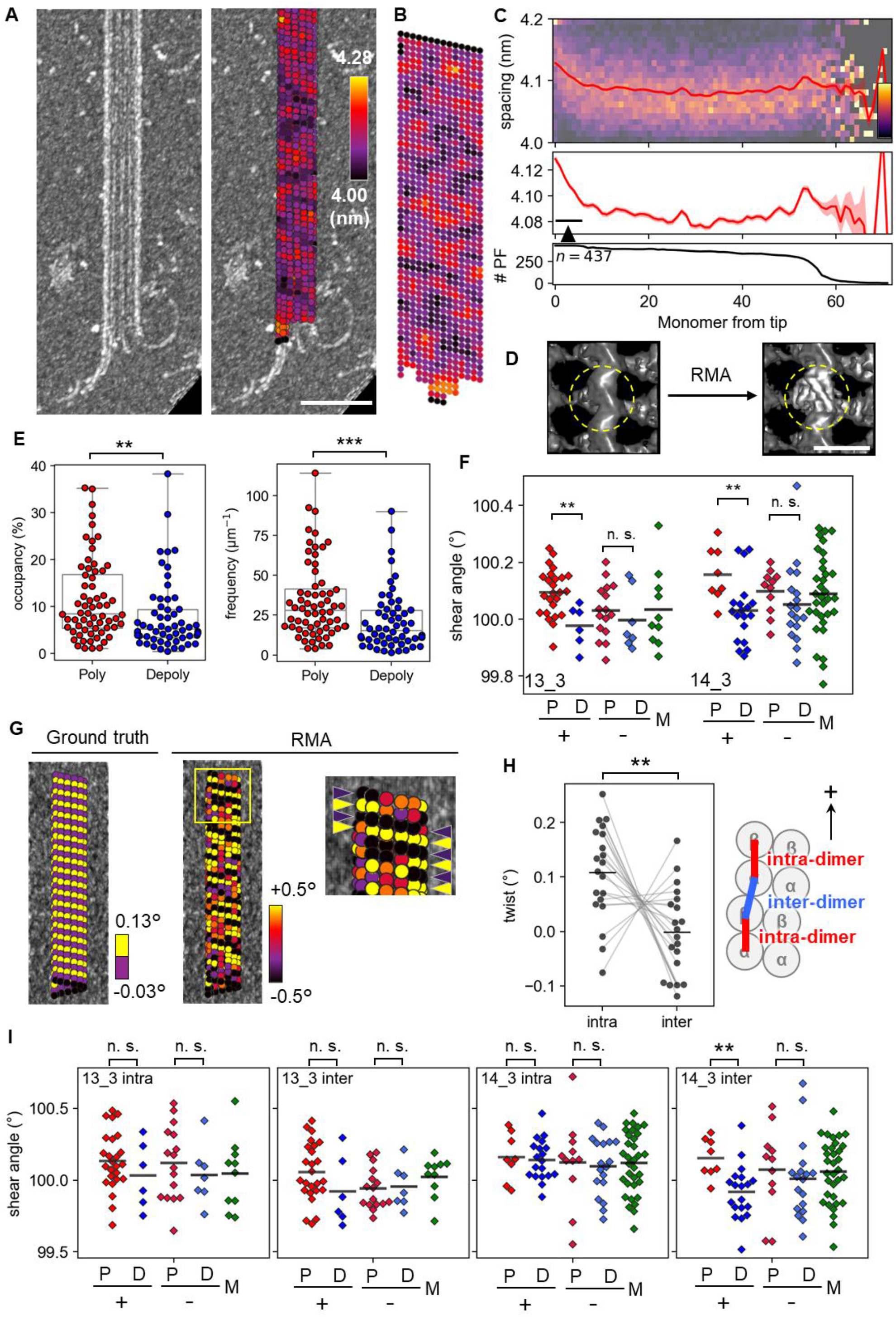
Lattice structure analysis of microtubule tips using RMA. (A) A representative estimation of lattice spacing of a polymerizing tip. Scale bar, 50 nm. (B) The flat view of the lattice spacing in (A). (C) A profile of lattice spacing from the very tips (top, middle) and the number of analyzed protofilaments (bottom). The heatmap indicates the distribution of the lattice spacing. The solid lines and the shaded regions are means ± S.E. The arrowhead indicates the relatively expanded tip region, which is less than eight dimers long. (D) Subtomogram averages of polymerizing 13_3 microtubule tips using tubulin coordinates before or after RMA alignment. The same threshold value was used to render both images. (E) Results of the lattice expansion analysis. Left, Occupancy of the expanded clusters. Right, Frequency of the appearance of expanded clusters. **: *p* < 0.01, ***: *p* < 0.001 (Mann-Whitney’s U test). (F) Plot of mean shear angles of each type of microtubule. Horizontal bars are the average of all dots. *: *p* < 0.05 (Welch’s t-test). (G) A representative simulation result of super twists. Tubulin molecules are colored by the twist angles. Estimated values were mean-filtered by five laterally connected tubulins. The zoomed image shows that changes in the twist angle were successfully detected (yellow and purple arrowheads). (H) Plot of the estimated super twists after ten simulations and a schematic illustration of the intra- and inter-dimer interface. *: *p* < 0.05 (t-test of related datasets). (I) Intra- and inter-dimer shear angle estimation using seam-searching and RMA alignment. **: *p* < 0.01 (Welch’s t test). n.s.: not significant.

### The combination of seam-searching and RMA alignment revealed interdimer shearing of polymerizing plus ends

Next, we conducted a real-space analysis of angular parameters using RMA. Unlike the estimation of spacing, the 0.05-nm resolution of the alignment methods is not sufficient for the estimation of local angular parameters; the positional change of a nearby tubulin by 0.1° lattice shearing, for example, is sin(0.1°) × 4.1 (nm) / 0.2615 (nm/pixel) = 0.027 pixels, which is far below the pixel size. However, we assumed that the per-microtubule average of shear angles still reflects the true microtubule lattice structures. As expected, the shear angle calculated from the molecule positions aligned by RMA successfully detected the difference of lattice shearing between the plus ends of polymerizing microtubules and the other ends for both 13_3 and 14_3 microtubules (Fig. 4F). These results indicate that RMA is applicable to estimating angular parameters.

Therefore, we attempted to determine the origin of the shear angle twist at the polymerizing plus ends using RMA. The shear angle can be altered by conformational changes within tubulin dimers and by the changes in the alignment between adjacent tubulin dimers. To confirm that RMA can detect supertwists at the intradimer and interdimer interfaces, we generated simulated microtubule tomogram images (Fig. 4G) simulating 200-nm microtubules with -0.03° twists in the intradimer interfaces and 0.13° twists in the interdimer interfaces (0.05° on average). The results show that RMA can detect the supertwist by comparing the twist angles averaged over intra- and inter-dimers (Fig. 4H). Therefore, if each tubulin can be correctly labeled with α or β, we can separately calculate shear angles for both tubulin populations. Although the microtubule seam makes it difficult to specify α/β-tubulins, previous works have demonstrated that an iterative trial of cross-correlation calculations can determine the correct seam location in 2D cryo-EM images (Zhang and Nogales, 2015) to successfully obtain high-resolution maps of undecorated microtubules (Zhang et al., 2018). We imitated this notion to implement the seam-search protocol for tomograms, in which all candidates of α- or β-tubulin average maps are calculated and the seam location is optimized by the tomogram correlation with α- and β-tubulin template images (Fig. S11A). We tested the accuracy of this protocol using kinesin-decorated microtubules (Fig. S11B– D) and found that our seam-search protocol can determine the seam location of a microtubule with <2-protofilaments precision using a 200-nm region of the microtubule. Using this protocol, we labeled all the aligned tubulin molecules with α or β. The success of this discrimination between α and β was confirmed by the presence and absence of a loop density specific for α-tubulin in the images obtained by averaging the subtomograms of each tubulin (Fig. S12). The ability to discriminate α/β-tubulins allowed us to separately evaluate the shear angles at inter- and intra-dimers in native microtubules. By comparing shear angles in microtubules during polymerization and depolymerization, only the shear angle at the inter-dimer in the 14_3 plus ends was significantly different, with a mean shear angle 0.24° larger during polymerization than during depolymerization. (Fig. 4I). This value was almost twice the increase in twist angle without considering supertwists (0.13°; Table S5), suggesting that inter-dimer shearing mostly contributed to overall lattice shearing. Although 13_3 microtubules followed the same trend, the sample size did not provide enough statistical power (1 - β = 0.26). Since GTP hydrolysis exclusively occurs in the inter-dimer interface, which causes a loss of γ-phosphate, this result implies that γ-phosphate only has a short-range effect on the microtubule lattice structure. Together, our results suggest that the lattice of microtubule polymerizing ends is sheared and that the lattice shearing is primarily caused by interdimer conformational changes.

### Constrained alignment revealed a preference for expansion to compaction in bending regions

Microtubules are not always straight, but often bend, especially within cells. Several MAPs are known to bind specifically to the bent regions of microtubules, so bending can be considered a physical cue for protein interactions (Bechstedt et al., 2014; Siahaan et al., 2022), but the detailed structure of bent microtubules has not been reported. In the bent region, individual protofilaments are assumed to be deformed differently and take different lattice parameters, making it difficult to assess. In the CFT analysis, we noticed that the estimation bias of the longitudinal lattice spacing by microtubule bending is larger in experiments (Fig. S6) than in simulations (Fig. S4F). We speculated that this difference is due to the expansion of the outer surface being larger than the compaction of the inner surface. To test this hypothesis, we analyzed the middle regions of the microtubules from the depolymerizing samples using RMA (Fig. 5A). We intentionally selected microtubules with various bends to cover a wide range of curvatures. The *η* value was introduced to label whether each tubulin molecule is inside or outside the bending (See Materials and Methods for its definition). As expected, a flat view of a strongly curved microtubule showed a clear correlation between *η* values and spacings (Fig. 5B, C). To quantitatively analyze the effect of bending, aligned tubulins were split into three groups: inner, side and outer regions (Fig. 5D), and the measured lattice spacings were plotted against the local curvatures. In the straight part of the microtubules, the three groups showed the same distribution of the lattice spacing. As the microtubule curvature increased, the increasing rate of the lattice spacing for the outer region was much larger than the decreasing rate of the inner region (Fig. 5E). Linear regression parameters indicated that the energy landscape of the microtubule expansion/compaction is biased; to achieve compaction, tubulins need ∼9 times more energy than for expansion for 13_3 microtubules and ∼13 times more for 14_3 microtubules (Table S6). We also analyzed the twist angle similarly, finding it was insensitive to the microtubule curvature (Fig. 5F; Table S6), consistent with the CFT analysis where shear angles were not affected (Fig. S6B). 15_3 microtubules also showed the same trend, although the sample size was small (Fig. 5E, F). Collectively, our analysis revealed that microtubule bending induces more expansion than compaction, while twist angles are not affected.

**Fig. 5:**
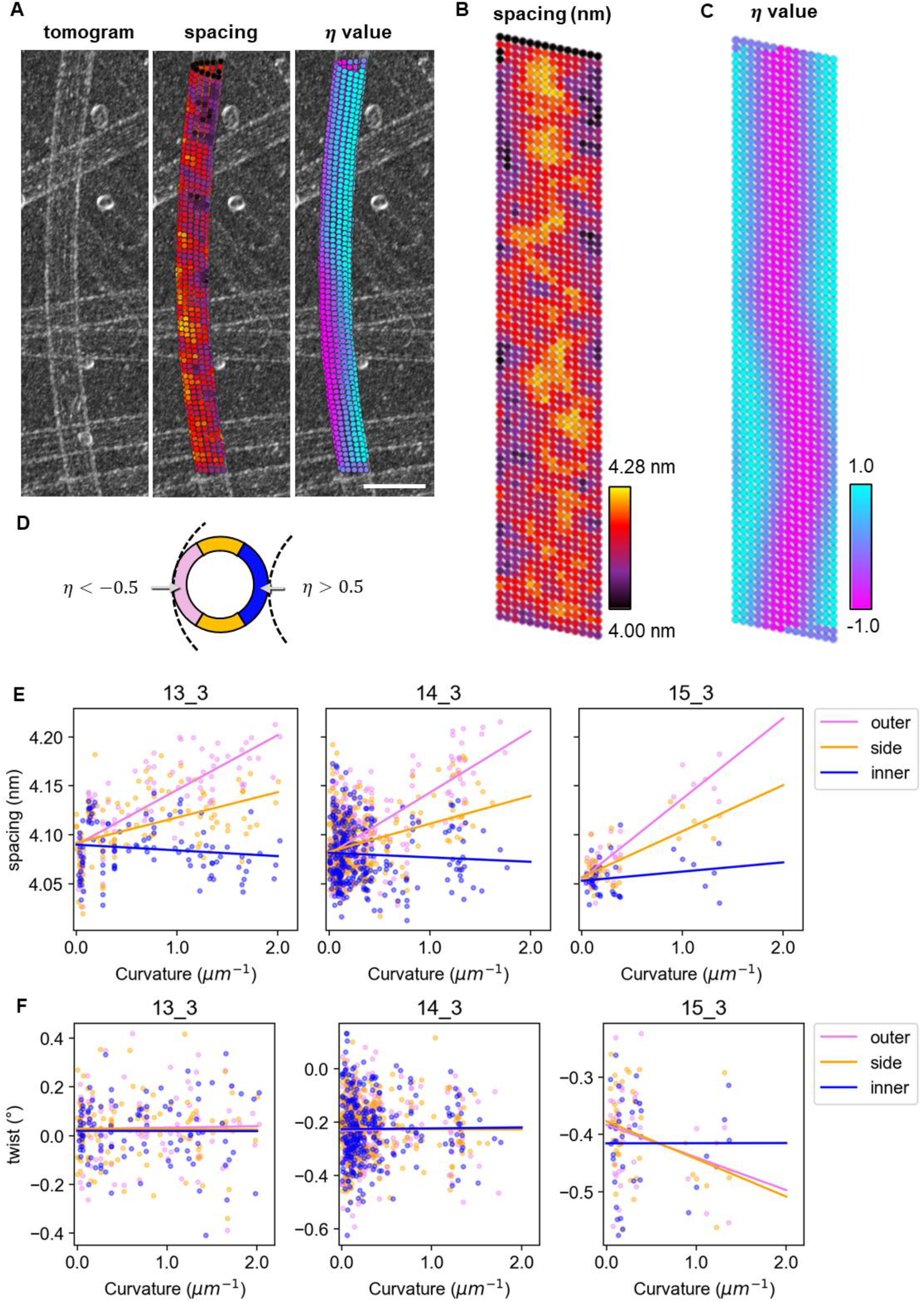
Curvature dependency of lattice parameters. (A) A representative filtered tomogram of a bent microtubule (left) with the molecules aligned by RMA. Molecules are colored by inter-molecular spacing (middle) or the *η* value (right). The *η* value was used to define the inside (*η* > 0) and outside (*η* < 0) of the bending. Scale bar, 50 nm. (B, C) Flat views of the molecules shown in A. Note that the molecules at the edges take *η* = 0 since they are out of the edges of the spline. (D) Definition of the inside, middle and outside of the bent microtubule. (E, F) Plot of local spacings (E) and twist angles (F) for each subregion (dots) and the result of the linear regression (lines). In (F), 13_3, 14_3 and 15_3 microtubules were analyzed separately.

In the present study, we developed new approaches to analyze the local structural properties of individual microtubules. Our analysis, which does not use averages, provides a deeper understanding of the local lattice structure of microtubules and its implications for polymerization, depolymerization, bending and the behavior of microtubule-associated proteins.

As for polymerization, our results on native polymerizing ends challenge previous assumptions. Based on the structures of artificially stabilized microtubules, polymerizing microtubule ends have been expected to have an expanded lattice (Alushin et al., 2014; LaFrance et al., 2022). However, our local-CFT and RMA analyses revealed that the lattice spacing of polymerizing ends is not extensively expanded but rather is closer to the compacted state observed in GDP-MTs (Fig. 2 and 5). Our observations are consistent with the behavior of MAPs that preferentially bind to the expanded lattice, such as KIF5C (Budaitis et al., 2022), TPX2^micro^ (Zhang et al., 2017) and the microtubule binding domain (MBD) of CAMSAP2/3 (Jiang et al., 2014), all of which do not track the polymerizing ends. Notably, the most distinctive feature of polymerizing microtubule plus ends was the change in the lattice shear angle (Fig. 2H, 5F), which was most likely caused by inter-dimer shearing (Fig. 4I). The fact that only the plus ends, which show a fast polymerization rate, showed a large inter-dimer shear angle may suggest that this angle correlates with the microtubule polymerization rate. The tip shearing probably contributes also to the difference in the off-axis fluctuation of protofilaments at the tips, as predicted by an atomistic molecular dynamics simulation of microtubule plus ends (Igaev and Grubmüller, 2022). Considering the GTPase activity of tubulin, lattice shearing is likely to correspond to the regions composed of GTP- or GDP•Pi-bound β-tubulins. The stochastic hydrolysis of GTP in the polymerizing ends should also contribute to the observed broad distribution of lattice spacings (Fig. 4E). Based on these results and assumptions, we propose that the sheared lattice works as a “cap” to promote polymerization and prevent the tip from catastrophe, similar to that traditionally ascribed to GTP caps (Fig. 6A).

**Fig. 6:**
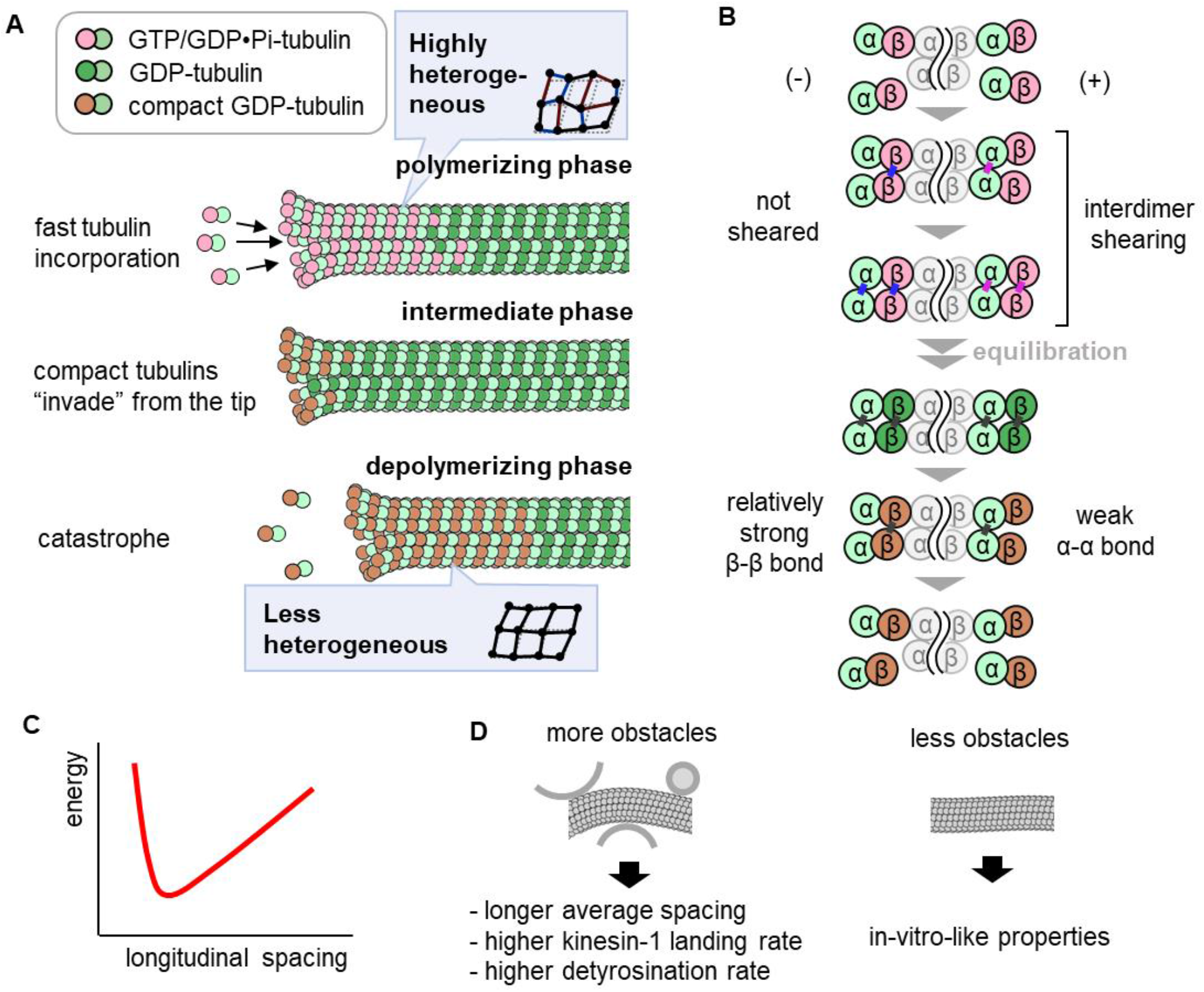
Heterogenic configuration of microtubules revealed by our non-averaging analysis. (A) Changes in microtubule tip structures during the polymerizing/depolymerizing cycle. β-tubulins are colored by their representative lattice structure, but they have different extents of structural heterogeneity, which could be caused by lattice fluctuations. (B) Molecular-level explanation of the difference between the plus and minus ends. Colors are the same as in (A). (C) A free energy landscape as a function of longitudinal spacing between tubulin monomers speculated from the RMA alignment results. (D) The effect of fluctuations on the mean expansion rate of microtubules.

How does the sheared lattice control microtubule dynamics? The share angle depends on the arrangement between adjacent tubulins, and this arrangement may affect the polymerization dynamics. Both longitudinal and lateral contacts between adjacent tubulins are required for microtubule formation since tubulins cannot stably exist as individual protofilaments nor as laterally connected short rings (Manka and Moore, 2018). In this perspective, changes in the lattice shear angle should affect the on- and off-rate of the tubulin at the corner of the lattice edges, resulting in changes in the balance between polymerization and depolymerization. An intriguing finding of the present study is the disparity in lattice structures between polymerizing plus and minus ends (Fig. 2H, 5F). A previous analysis of the averaged structures of doublecortin-bound GDP-Pi-MT and GDP-MT demonstrated significant changes in the interface of the α-α and β-β lateral contacts upon phosphate release (Manka and Moore, 2018), which likely serves as the primary cause of lattice shear. So far, the residues responsible for the difference between the two tips have not been identified, as the averaged structure of doublecortin-bound GDP-Pi-MT consists of a mixture of plus- and minus-end microtubules. However, it is plausible that the configuration of soluble tubulin dimers determines the optimal shear angles of the plus and minus ends (Fig. 6B). During polymerization, tubulin dimers exhibit curvature at the intradimer interface before straightening into a cylinder (Gudimchuk and McIntosh, 2021). As a tubulin dimer approaches the exposed β-tubulins at the plus end, the distance between laterally adjacent α-tubulins is smaller than that between β-tubulins. As a result, the first lateral bond is expected to be formed at the α-α interface in the plus ends, whereas it occurs at the β-β interface in the minus end. A difference in the optimal shear angle between α-α and β-β will result in different lattice parameters. Therefore, the difference in the order of the α-α and β-β contact formation can explain the differences between the two types of microtubule tips during polymerization.

Consistent with the proposed model, the depolymerizing tips had smaller shear angles than the polymerizing tips (Fig. 2H, 2I). However, both plus and minus ends of depolymerizing microtubules showed similar shear angles (Fig. 2, 5). Therefore, to elucidate the molecular basis of the difference in the tubulin off-rate between the two ends (Strothman et al., 2019), we must await future molecular dynamics simulations that incorporate the lattice information revealed in the present study. Nevertheless, the aforementioned differences in the order of breakage between α-α and β-β bonds may contribute to the off-rate variation. In addition, the depolymerizing tips showed characteristic values not only in shear angle but also in lattice spacing; the lattice spacing at the depolymerizing tips was shorter than that in the middle regions of GDP-MT (Fig. 2G), suggesting that depolymerization is preceded by lattice compaction (Fig. 2G). This lattice compaction may also be related to the smaller lattice fluctuation of depolymerizing ends (Fig. 4E). The compacted lattice spacing and the rare appearance of expanded clusters on depolymerizing microtubules are consistent with existing evidence suggesting that lattice expansion typically inhibits depolymerization (Alushin et al., 2014; Morikawa et al., 2015; Liu and Shima, 2023).

Using RMA, we found that microtubule bending leads to greater expansion in the outer protofilaments compared to compaction in the inner protofilaments while having a minimal effect on the twist angle (Fig. 5E, F). This observation suggests a biased free energy landscape for microtubule expansion/compaction at the bending regions (Fig. 6C), which in turn implies that the average microtubule lattice spacing should be larger in confined spaces, such as the cytoplasm, where the presence of obstacles causes microtubules to bend (Brangwynne et al., 2007), compared to in vitro experiments, where microtubules are typically immobilized on a larger glass surface (Fig. 6D). As a result, MAPs with a higher affinity for expanded lattices are more likely to be recruited to the curved regions of microtubules in cells (Zhang et al., 2017; Shima et al., 2018). Changes in kinetic parameters of MAPs by bending have also been observed with high-speed atomic force microscopy: kinesin has a higher on-rate and lower off-rate for bent regions of microtubules, leading to slower translocation (Nasrin et al., 2021). Although the molecular dynamics simulation in that article suggested that both lattice expansion and compaction lead to stronger kinesin-microtubule interactions, it is important to note that the interactions may be enhanced merely by the lattice spacing biasing to the expanded state (Nakata et al., 2011; Shima et al., 2018). Furthermore, it has been reported that bending can influence the post-translational modifications (PTMs) of microtubules (Janke, 2014). In particular, recent research on microtubule PTMs has shown that the activity of VASH1/SVBP, an enzyme responsible for microtubule detyrosination, is regulated by microtubule lattice expansion (Yue et al., 2023). This finding raises the intriguing possibility that microtubule detyrosination can be affected by physical bending, thereby elucidating the mechanism by which transient deformation of the microtubule shape leads to long-term chemical modifications. The interplay between microtubule shape, lattice structure, modifications and MAP interactions provides a promising framework for comprehensively understanding the intricate behavior of the microtubule network.

Besides the microtubule curvature, the distinction between polymerizing plus and minus ends provides valuable structural cues for MAPs. For instance, KIF1A, a kinesin motor protein involved in synaptic transport in axons, dissociates more frequently from dynamic plus ends, and the loop 11 region of KIF1A has been shown to be essential for plus end recognition (Guedes-Dias et al., 2019). Although loop 11 binds to the intra-dimer interface of a tubulin dimer, a previous study suggested that loop 11 mediates mechanical forces through its loop 8 region to H11 of α-tubulin, which sits close to the inter-dimer interface (Shima et al., 2018). Therefore, KIF1A may sense the inter-dimer shear angle through the mechanical relay to distinguish dynamic plus ends. In contrast, microtubule end-binding proteins (EBs) track both the plus and minus ends of polymerizing microtubules. The insensitivity of EBs to the plus and minus ends indicates that the tip-tracking mechanism is not merely dependent on the change in the angular parameters of the microtubule lattice, a characteristic conformational change of EB-bound microtubules (Zhang et al., 2015). Although EBs bind to the inter-dimer interface, which is suitable for sensing the lattice shearing of polymerizing plus ends (Fig. 4I), amino acid residues that are important for end-tracking are located mostly in the interface with β-tubulins (Maurer et al., 2012). This interaction is consistent with our idea that EB-binding is insensitive to lattice shearing, as shearing can occur without changing the interface between EBs and β-tubulins. Therefore, EBs recognize microtubule polymerizing ends either by direct interactions with GTP γ-phosphate or by sensing small conformational changes in β-tubulins induced by the γ-phosphate, changes that do not affect the overall lattice parameters.

A persistent challenge that remains unresolved in the present study is the distribution of GTP γ-phosphate and the associated Mg^2+^ ions. Comparisons of microtubule average structures under different nucleotide and binding protein conditions suggest the potential absence of either or both γ-phosphate and Mg^2+^ during GTP hydrolysis (Zhang et al., 2015). Elucidating the relationship between the lattice structures (Figure 6A) and the presence or absence of γ-phosphate and/or Mg^2+^ is a critical step in unraveling the molecular mechanisms underlying microtubule dynamic instability. Unfortunately, extracting such high-frequency information from small microtubule fragments is quite challenging. In addition, the relatively lower resolution of cryo-ET compared to single-particle cryo-EM analysis hinders the direct detection of γ-phosphates even with subtomogram averaging techniques (Fig. S12). Advances in image analysis workflows and experimental methods are essential to overcome these limitations.

The present study has introduced innovative non-averaging methods for the structural characterization of individual microtubule local lattices. These methods have significant potential as powerful tools for investigating local and/or transient interactions of microtubules with MAPs and how such interactions regulate specific cellular events. Furthermore, numerous biological structures exhibit similar cylindrical periodicity as microtubules and can change their structure under different biological circumstances, such as actin filaments (McGouph et al., 1997), tau filaments (Fitzpatrick et al., 2017) and amyloid fibrils (Tycko, 2015). Our methods will help to study such polymorphisms of cylindrical structures. Furthermore, although these methods were developed for non-averaging analysis, they can also contribute to improving the quality of averaged images of subtomograms. This is because Viterbi alignment and RMA can prevent the misalignment of electron densities, which leads to poor image quality, by accurately estimating the arrangement of the constituent molecules in the cylindrical structures. This feature is particularly promising when dealing with complex and noisy cellular environments, as these methods are more tolerant of noise than previous alignment algorithms. For these reasons, we believe our methods will make a significant contribution to future structural biology.

## STAR★Methods

### Key resources table

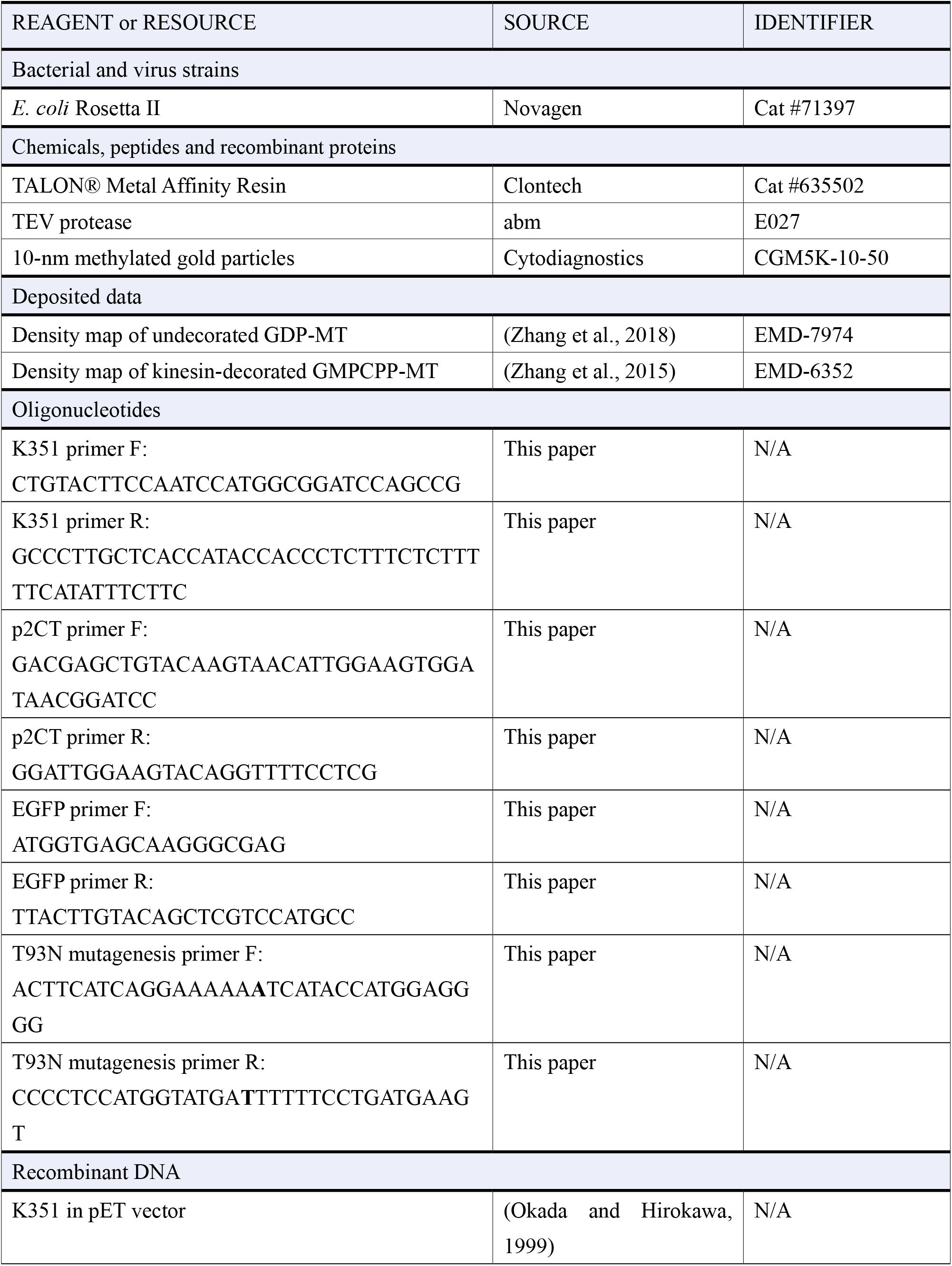
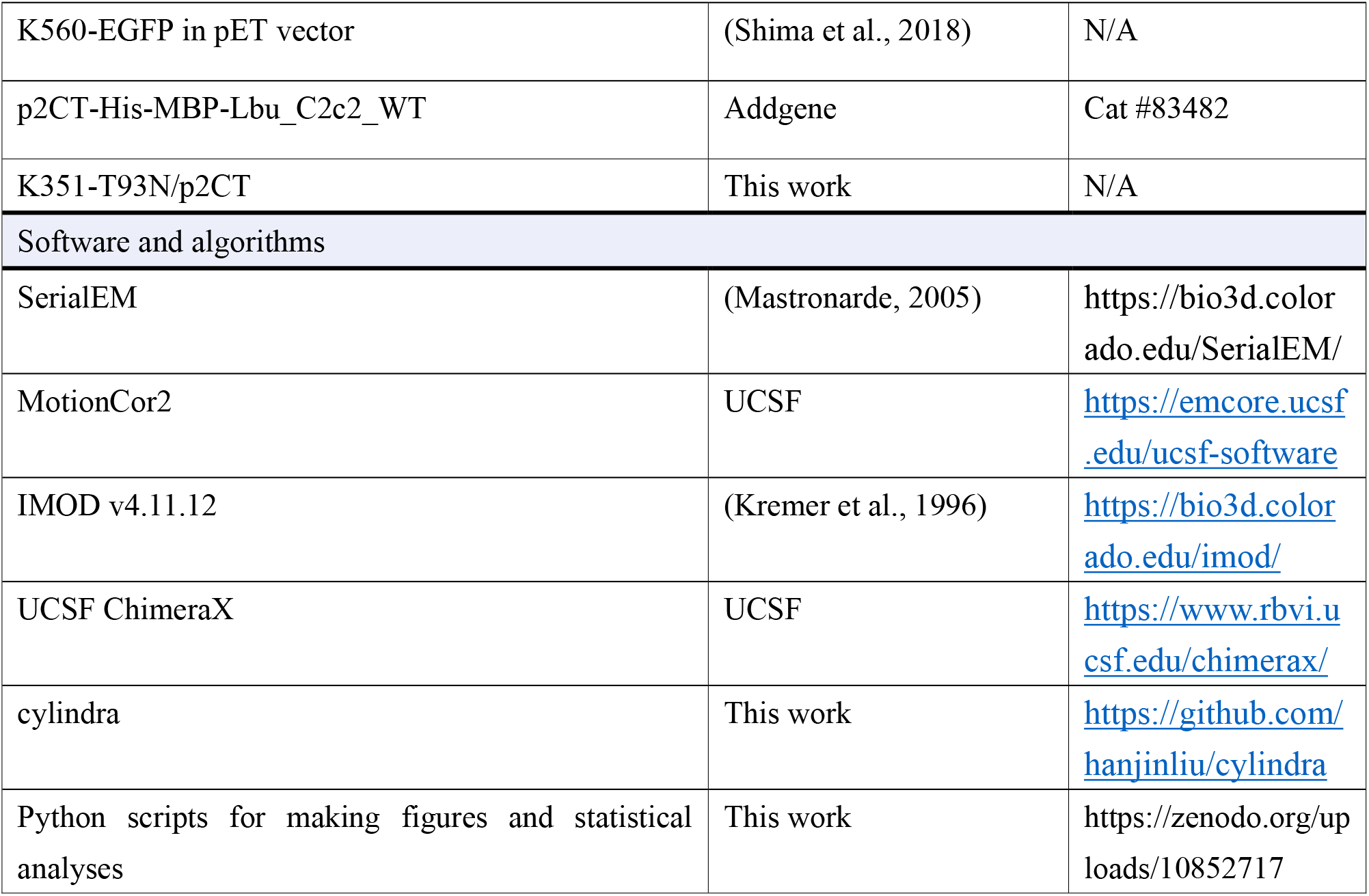

### Resource availability

#### Lead contact

Further information and requests for resources and reagents should be directed to and will be fulfilled by the lead contact, Tomohiro Shima.

#### Material availability

Plasmids generated in this study will be shared upon request.

#### Data and code availability

All image data sets will be provided upon request. Functions, graphical user interfaces and scripts for the simulations are available on GitHub (https://github.com/hanjinliu/cylindra). Scripts to reproduce all the analysis workflows are deposited on Zenodo (https://zenodo.org/uploads/10852717).

## Method details

### Protein purification

Tubulin was purified from porcine brains by three cycles of polymerization and depolymerization (Castoldi and Popov, 2003), dissolved in PEM buffer (100 mM PIPES, 1 mM EGTA and 2 mM MgSO_4_, pH 6.9 adjusted with KOH) or BRB80 buffer (80 mM PIPES, 1 mM EGTA and 2 mM MgSO_4_, pH 7.3 adjusted with KOH) supplemented with 0.1 mM GTP and stored in liquid nitrogen until use.

Rigor kinesin-1 mutant protein (residues 1 – 351 of mouse KIF5C with the T93N mutation) was purified from *E. coli*. Kinesin-1 gene was cloned into p2CT vector with 6xHis-tag maltose binding protein (6xHis-MBP, the solubilization tag) connected with the TEV protease recognition sequence at the N-terminus and EGFP at the C-terminus, and transformed into Rosetta II competent cells (71397; Novagen). Cells were cultured in 2x YT medium (16 g/L bacto-tryptone, 10 g/L yeast extract and 5 g/L NaCl) supplemented with 10 μg/mL ampicillin until the OD600 reached ∼0.5. 0.5 mM of isopropyl β-D-thiogalactopyranoside (IPTG) was added to the medium and culture at 24°C for 5.6 hours to induce protein expression. The bacteria pellet was resuspended in lysis buffer (50 mM Tris, 2 mM MgCl_2_, 250 mM NaCl and 5 mM imidazole, pH 8.0), supplemented with protease inhibitor cocktail (1 mM 4-(2-aminoethyl) benzenesulfonil fluoride, 20 μM Leupeptin, 8.7 μM Pepstatin and 1 mM N-p-tosyl-L-argininie-methyl ester), sonicated and pelleted by centrifugation. The supernatant was incubated with TALON® Metal Affinity Resin (Clontech, #635502) at 4°C for 30 minutes and the resin was washed with six column volumes of wash buffer A (50 mM Tris, 2 mM MgCl_2_, 250 mM NaCl and 10 mM imidazole, pH 8.0) and six column volumes of wash buffer B (50 mM Tris, 2 mM MgCl_2_, 250 mM NaCl and 20 mM imidazole, pH 8.0). Rigor kinesin-1 was eluted with four column volumes of elution buffer (50 mM Tris, 2 mM MgCl_2_, 250 mM NaCl and 200 mM imidazole, pH 8.0). To cleave the N-terminal 6xHis-MBP, 0.17 U/μL of TEV protease (abm, E027) was added to the elution and dialyzed overnight in the dialysis buffer (20 mM PIPES, pH 6.7) supplemented with 1 mM Tris(2-carboxyethyl)phosphine hydrochloride (TCEP). After separating the cleaved 6xHis-MBP by mixing with TALON® Metal Affinity Resin, rigor kinesin-1 was concentrated with a 30-kDa-cutoff centrifugal filter (UFC503096; Merck) and stored at -80°C.

### Cryo-ET grid preparation

For electron tomography of non-dynamic GDP-MT, 40 μM tubulin solution was supplemented with 1 mM GTP and cleaned by centrifugation (20,000 × g, 4°C, 2 min) before polymerization at 37°C. After 50 minutes of polymerization, non-polymerized tubulin was removed by ultra-centrifugation (436,000 × g, 35°C, 5 min). Microtubules were stored as pellets and resuspended in 20 μM tubulin solution with BRB80 buffer and 1 mM GTP to adjust the microtubule concentration. Microtubules were mixed with 10-nm methylated gold particles (Cytodiagnostics CGM5K-10-50) to produce a fiducial marker along with 0.01% NP-40 and 10 μM tubulin solution with BRB80 buffer and 1 mM GTP before the solution was dropped onto the grids (QUANTIFOIL R 1.2/1.3 Cu 200). The grids were blotted with vitrobot (Thermo Fisher Scientific; 22°C, 100% humidity, blot force 15 for 6 seconds), vitrified with liquid ethane and stored in liquid nitrogen until observation.

For GMPCPP-MT, 5 μM tubulin solution was supplemented with 0.1 mM GMPCPP and cleaned by centrifugation (20,000 × g, 4°C, 2 min). After 20 min of nucleation at 27°C and 50 minutes of polymerization, non-polymerized tubulin was removed by ultra-centrifugation (300,000 × g, 35°C, 7 min). Microtubules were stored as pellets and resuspended in BRB80 buffer to adjust the microtubule concentration. 0.2 μM GMPCPP-MT (assuming 70% of the tubulin was polymerized) was first dropped onto the grid and subsequently exchanged to kinesin mixture containing 0.5 μM rigor kinesin-1, 0.01% NP-40 and methylated gold particles in BRB80 buffer. This grid was incubated in vitrobot for 5 min and blotted in the same protocol as GDP-MT.

For electron tomography of polymerizing microtubules, tubulin was concentrated by an additional cycle of polymerization/depolymerization and dissolved in PEM buffer supplemented with 1 mM GTP. This tubulin sample was cleaned by centrifugation (20,000 × g, 4°C, 2 min) and mixed with gold particles as a fiducial marker and 0.01% NP-40 before polymerization. To minimize the possibility of including depolymerizing ends and to maximize the size of GTP caps, polymerization was conducted with 48 μM of tubulin at 37°C for 3 or 8 minutes. After dropping the microtubule solution onto the grids, which took ∼1 min at room temperature, the grids were incubated at 37°C and 100% humidity in vitrobot for 1 min (polymerized for 5 or 10 minutes in total). The grids were blotted at blot force 15 for 6 seconds, vitrified with liquid ethane and stored in liquid nitrogen until observation.

For electron tomography of depolymerizing microtubules, 40 μM tubulin was supplemented with 1 mM GTP and polymerized for 30 minutes at 37°C. To remove non-polymerized tubulin molecules, polymerized microtubules were purified by ultra-centrifugation (300,000 × g, 35°C, 7 min). Microtubule pellets were resuspended in 25% PEG200 and 3/4× BRB80 buffer and diluted by 10 times in 20 μM tubulin solution with 1 mM GTP just before adding onto the grids. To start depolymerization, the grids were blotted at blot force 15 for 3 seconds, and tubulin-free buffer (0% or 3% PEG200, gold particles, 0.01% NP-40 in BRB80 buffer) was added onto the grid inside the vitrobot. Immediately after, the same freezing protocol as for polymerizing microtubules was applied.

### Tomogram Data Collection and Reconstruction

Tilt series were acquired using a Titan Krios G3 (ThermoFisher Scientific) at 300 kV with a K3 detector in the counting mode and a Quantum-LS with a slit width of 30 eV. Images were acquired at 33,000x magnification (pixel size ∼0.26 nm) with defocus between -2 and -5 μm and final electron dose between 100 and 110 e^-^/Å^2^, and each image was subdivided into 10-20 movie frames. The range of tilt angles was set between -60° and 60° at 2° increments. All data collections were operated on a SerialEM.

All acquired image frames were corrected using MotionCor2 (Zheng et al., 2017). Image processing, including tilt series alignment, contrast transfer function (CTF) correction, gold bead erasure and weighted back projection, were conducted using the etomo interface of IMOD (v4.11.12) and saved as cropped 32-bit floating images.

### Tomogram Simulation

To reconcile the experimental procedures of the tomogram acquisition as much as possible, we first created microtubule models by placing tubulin density maps made from the map of undecorated GDP-MT (EMD-7974; Zhang et al., 2018) at arbitrary coordinates. The model was projected with view directions ranging from -60° to 60° at 2° increments to make a noise-free tilt series. White Gaussian noise was added to the tilt series before the weighted back-projection (inverse Radon transformation).

The standard deviation of the Gaussian noise equivalent to the experimental tomograms (*σ*_*exp*_) was determined as follows. Tomograms containing a 200-nm-long microtubule were simulated with standard deviations *σ* = 3.0, 3.5, 4.0-, while the maximum intensity of the template αβ-tubulin density map was normalized to 1. The subtomogram alignment of 600 tubulins was conducted for each tomogram, and the mean intensity of the 5.8×5.8×5.8 nm^3^ average image (single tubulin molecule) was defined as the “signal”. The average image was subtracted from the subtomograms to create noise-only images, whose standard deviation (*n*(*σ*)) was calculated as the “noise”. The relationship between *σ* and the noise-to-signal ratio *n*(*σ*)/*s* can be considered the calibration curve between the noise in the tilt series and that in the reconstructed tomograms. Six polymerizing microtubules and three depolymerizing microtubules from three independent cryo-EM grids were selected from the reconstructed tomograms of polymerizing microtubules; *s*(*σ*_*exp*_) and *n*(*σ*_*exp*_) were calculated in the same way, and *σ*_*exp*_ was regressed by the calibration curve *σ* vs. *n*(*σ*)/*s*.

### Cylindric Fourier Transformation (CFT)

Microtubule fragments (for local-CFT) or straightened microtubules (for glocal-CFT) were transformed from the Cartesian coordinate system (*x*_*c*_, *y*_*c*_, *z*_*c*_) to the cylindrical coordinate system (*r, y, θ*) to crop out tubulin density in a sheet-like arrangement. For example, if the microtubule is parallel to the *y*_*c*_ axis, the coordinate system becomes:

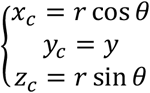

3D images of a microtubule sheet fragment, whose dimension is composed of a radial axis (*r*), longitudinal axis (*y*) and angular axis (θ), were analyzed by a local discrete Fourier transformation with (1, α_*y*_, α_θ_)-fold up-sampling:

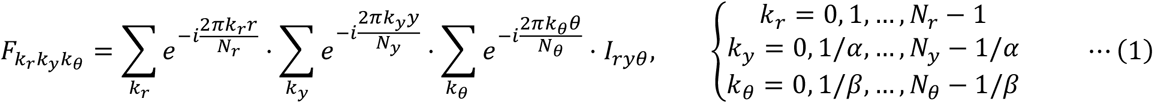

where *I*_*ry*θ_ is the image in cylindrical coordinates, *N*_*X*_ is the size of the image in the *x*_c_ axis and *i* is the imaginary unit. To avoid unnecessary calculations, frequency domains around the coordinates corresponding to 4-nm periodicity in the *r* -direction and 2 *π* /13 radian periodicity in the θ -direction were extracted. The cylindric power spectrum 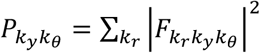was used to detect the peak positions of the longitudinal periodicity 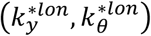 and lateral periodicity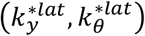. The lattice helical pitch *λ*_y_, rise angle θ_*r*_, protofilament number *N*_*PF*_ and skew angle θ_*s*_ can be directly calculated from these peak positions using the following formulae:

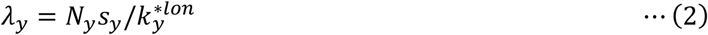

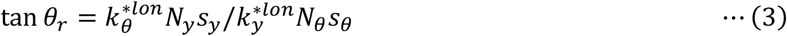

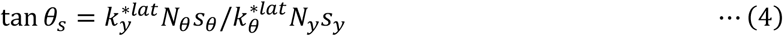

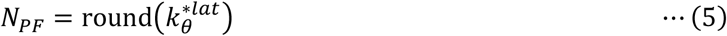

where *s*_*X*_ is the image scale factor (nm/pixel) of the *x*_*c*_ axis of the cylindric image. Note that *θ*_*r*_ and *θ*_*s*_ are defined on the cylinder surface so that they are distorted in the Cartesian coordinate system. Lattice spacing *L*_*y*_, rise length *L*_*r*_, twist angle *θ*_*tw*_ and start number *S* are calculated as follows:

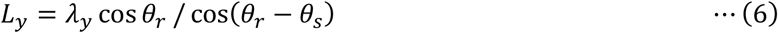

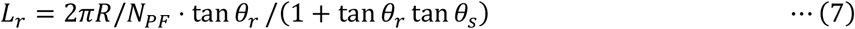

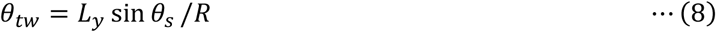

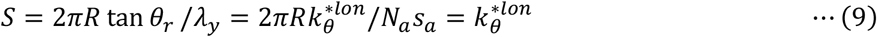

where *R* is the radius corresponding to the center of the *r* axis of the cylindric image. Note that *N*_*θ*_ is proportional to *R*, so that tan *θ*_*r*_ *∝ R* and tan *θ*_*t*_ *∝* 1/*R. λ*_*y*_, *N*_*PF*_, *L*_*r*_ and *S* are independent of *R*. The lattice shear angle *θ*_*sh*_ is calculated as follow:

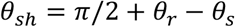

### Spline Fitting and CFT Analysis

Microtubules were first fitted to spline curves by rough spline fitting and further refinement using subtomogram alignment of the tubulin molecules as follows. Microtubule center curves were manually drawn, and sub-volumes containing microtubule fragments were sampled at 49-nm intervals. For each sub-volume, the center of the microtubule fragment was estimated using a mirrored auto-correlation in 3D (a 2D version was introduced in Blestel et al., 2009). Spline curves were updated using the newly estimated centers of all microtubule fragments (rough spline fitting). The mean radius of each microtubule was estimated by the radial profile from the spline, which was used to build the cylindric coordinate system surrounding the spline. The global-CFT of each microtubule was calculated to measure the mean lattice parameters, which were used to initialize all tubulin coordinates. The tubulin coordinates were shifted to the center of the corresponding tubulin molecules by subtomogram alignment with a search space of 0.8 nm x 0.8 nm x 0.8 nm using a zero-normalized cross-correlation with a template image of a αβα-tubulin trimer created from the map of an undecorated GDP-MT (EMD-7974; Zhang et al., 2018). The shifted coordinates were used to draw spline curves that precisely pass through the center of the microtubules.

After a microtubule is fitted to a spline curve, the first derivatives 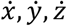 and the second derivatives 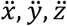 can be calculated at any points on the spline. These values are used to estimate the local curvature *κ* of the spline based on the following formula:

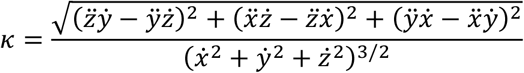

Local radii of 49-nm fragments were estimated by the radial profile of the spline. To avoid radius-sensitive lattice parameters being affected by the difference in the distribution, the mean radius for each protofilament number was calculated from the local radii and used as the consensus radius to build the final cylindric coordinate system along the splines.

For the local-CFT analysis of GDP-MT and GMPCPP-MT in Fig. 1, 49-nm-long sub-volumes of microtubule fragments were sampled at 49-nm intervals from the image binned by 2x2x2 (resulting in ∼0.5 nm/pixel). For the local-CFT analysis of polymerizing and depolymerizing microtubule tips, sub-volumes were sampled similarly, but the sampling interval was set to 8.16 nm. The global-CFT analysis was conducted for every microtubule to determine microtubule N_S structures. The results of the local- and global-CFT analysis were exported as parquet files for further analysis and plotting.

### Constrained Subtomogram Alignment

Prior to running Viterbi alignment or RMA, the coordinates and orientations of tubulin molecules were initialized using the lattice parameters determined by global-CFT. For each molecule, a 0.8 nm x 0.8 nm x 0.8 nm sub-volume was sampled by cubic interpolation, and the cross-correlation landscape *C*_*i*_ was calculated using a template image created from the high-resolution density map of undecorated GDP-MTs (EMD-7974). This landscape was used to calculate the energy of a microtubule given a set of tubulin coordinates {***x***_***i***_ | *i* = 0, 1, … }:

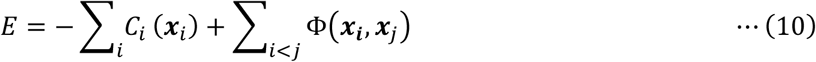

where Φ is the “binding energy” function.

In Viterbi alignment, only longitudinal connections are considered. Molecules are split into groups according to which protofilament they belong to, and the energy for the *k*-th protofilament:

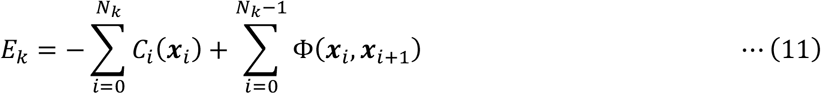

is the target for minimization. Two types of binding energies are considered in Viterbi alignment: the energy of the longitudinal distance *τ*_|_ and the energy of the longitudinal angle relative to the tubulin orientation *τ*_*∠*_

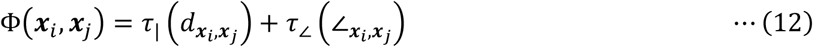

where 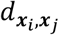 is the Euclid distance between ***x***_*i*_ and ***x***_*j*_, and 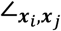 is the angle between the vector ***x***_*i*_ − ***x***_*j*_ and the tubulin orientation corresponding to the longitudinal axis of the microtubule. The infinite potential well is used for *τ*_|_ and *τ*_*∠*_

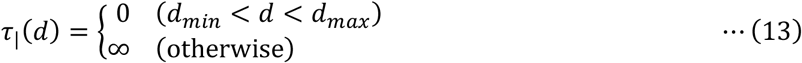

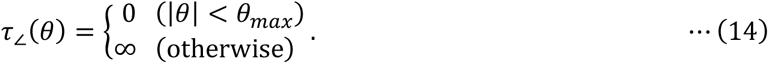

Compared to the Viterbi algorithm used in the hidden Markov model, −*C*_*i*_(***x***_*i*_) corresponds to the emission probability from state ***x***_*i*_ and Φ(***x***_*i*_, ***x***_*i*+1_) to the transition matrix, with slight differences such as the transition matrix not being a constant matrix throughout the data series. Using this analogy, the global minimization of *E*_*k*_ is achieved by the Viterbi algorithm (Viterbi, 1967).

In RMA, tubulin coordinates are determined using a random process based on simulated annealing. Simulated annealing is an iterative method to optimize parameters that minimize energy *E*. The parameters are updated by random walking inside the parameter space, while the transition probability from a parameter set to another that causes the energy change *ΔE* is defined by the Boltzmann distribution *exp*(−*ΔE*/*T*), where *T* is the temperature. By cooling down during the parameter updates, the energy will converge to the minimum. In RMA, the energy of the lateral distance *τ*_−_ is also considered in combination with *τ*_|_ and *τ*_*∠*_, and the trapezoidal potential is used instead of the infinite potential well:

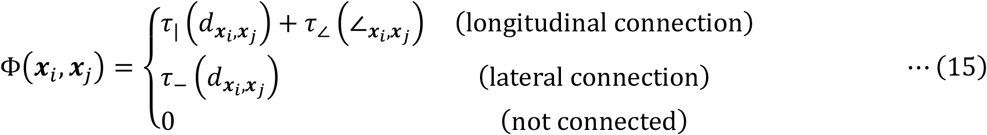

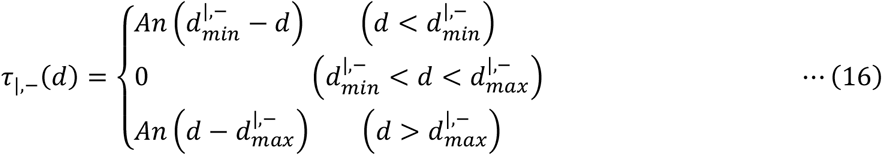

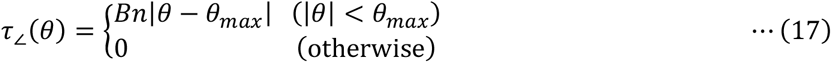

where *n* is the number of annealing steps, and *A, B* are positive constant values throughout the annealing. The trapezoidal potential starts as a flat potential (*n* = 0), but during annealing, the boundaries get steeper and the potential eventually becomes an infinite potential well (*n →* ∞).

Viterbi alignment and RMA are computationally intensive, so that they were implemented in Rust (v1.69.0) and compiled with maturin (v0.14.17) to be called in Python runtime via PyO3 (v0.19.0).

### Post-alignment Lattice Parameter Calculation

Let the spline curve of length *L* be ***C***(*s*) ∈ ℝ^3^ (0 ≤ *s* ≤ *L*) and *s*_*i*_ be the spline coordinate (distance along the spline from the starting point) corresponding to the *i* -th tubulin molecule at ***r***_*i*_ in the world coordinate. The lattice parameter calculation begins by converting any vector ***v*** starting from ***r***_*i*_ in the world coordinate and *s*_*i*_ in the spline coordinate into cylindric coordinate representation (*v*_*r*_, *v*_*y*_, *v*_*θ*_) (Fig. S8). Normal vectors of a cylinder surrounding the spline at point ***r*** and corresponding spline coordinate *s* can be calculated as

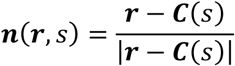

and the spline tangent unit vector at coordinate *s* can be calculated as

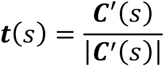

Changes in spline coordinate *Δs* caused by vector ***v*** can be estimated by projection to the spline:

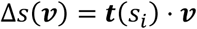

and *v*_*r*_, *v*_*y*_ are calculated by projection to the surface vector and tangent vector at the midpoint *s*_*i*_ + *Δs*(***v***)/2 (Fig. S8B):

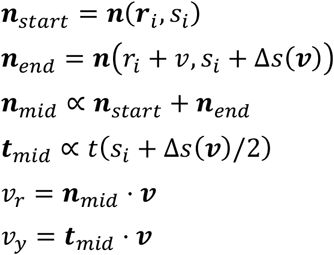

*v*_*θ*_ is, however, a nonlinear projection to the cylinder surface. Therefore, it is geometrically calculated by surface vectors and the cylinder radius *R* (Fig. S8C):

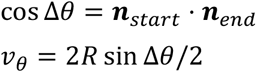

Vectors between longitudinally or laterally adjacent tubulin molecules ***v***^*lon*/*lat*^ were used to calculate the lattice spacing *L*_*y*_, skew angle *θ*_*s*_, rise angle *θ*_*r*_ and shear angle *θ*_*sh*_ :

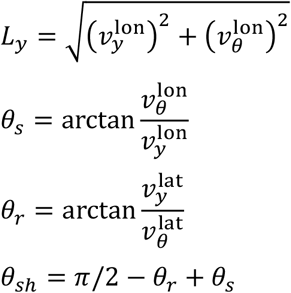

### Relationship between Microtubule Curvature and Structure

Given a spline curve ***C***(*u*) ∈ ℝ^3^ (0 ≤ *u* ≤ 1) and *i*-th molecule at ***r***_*i*_ in the world coordinate and *u*_*i*_ in the spline coordinate, the second derivative of spline ***C***^′′^(*u*) points inward on the curve (same as the acceleration vector of any curve). Therefore, the angle *η*_*i*_ between ***C***^′′^(*u*_*i*_) and the spline-to-molecule vector ***r***_*i*_ − ***C***(*u*_*i*_) can be used as an indicator of whether a molecule locates on the inner side of a curve.

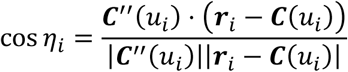

0° < |*η*_*i*_| < 60°, 60° < |*η*_*i*_| < 120°, 120° < |*η*_*i*_| < 180° were defined as the inner, side and outer region of a curve, respectively. The local curvature at *u*_*i*_ is used for the curvature of the microtubule region that the *i*-th molecule belongs to. Molecules were split into 49-nm pieces, and the curvatures and the longitudinal intervals were averaged within each piece for plotting and linear regression.

### Microtubule Seam-search

To determine the microtubule seam location without using associated proteins as the fiducial, we modified the seam-search protocol for 2D images (Zhang and Nogales, 2015) to implement this protocol for tomograms. Tubulin positions and orientations were first determined by alignment using RMA. All possible assignments of α/β-tubulin (twice the protofilament number) were tried for each microtubule, and the subtomogram averages around α- and β-tubulin were calculated. The cross-correlation between the averages and the α- or β-tubulin monomer template image were calculated, and the difference in the correlation was plotted against the seam location. The seam location that maximizes the value (assuming an average image of β-tubulin fits best and that of α-tubulin fits worst) was used as the optimal estimation.

To validate this seam-search protocol, tomograms of kinesin-decorated GMPCPP-MT were used. The seam location was determined precisely using the correlation between kinesin fiducials in the tomogram and the kinesin template image made from kinesin-decorated GMPCPP-MT (EMD-6352).

## Quantification and statistical analysis

All the quantification and statistical analyses were conducted using Python. The scripts are uploaded to Zenodo (https://zenodo.org/uploads/10852717).

## Acknowledgement

We thank the members of the Uemura and Kikkawa laboratory for valuable discussions. This work is supported by JSPS KAKENHI (23KJ0472 to H.L.; 18K06147, 19H05379, and 21H00387 to T.S.) and Basis for Supporting Innovative Drug Discovery and Life Science Research (BINDS) from AMED under Grant Number JP21am0101115 (support number 3013).

## Author Contributions

Conceptualization, H.L. and T.S.; Data curation, H.L. and H.Y.; Methodology, H.L., H.Y., M.K. and T.S.; Investigation, H.L. and H.Y.; Analysis, H.L. and H.Y.; Supervision, M.K. and T.S.; Visualization, H.L.; Writing –original draft, H.L.; Writing –review & editing, H.L., H.Y., M.K., and T.S.

## Supplementary Information

### Supplementary Figures

**Fig. S1:**
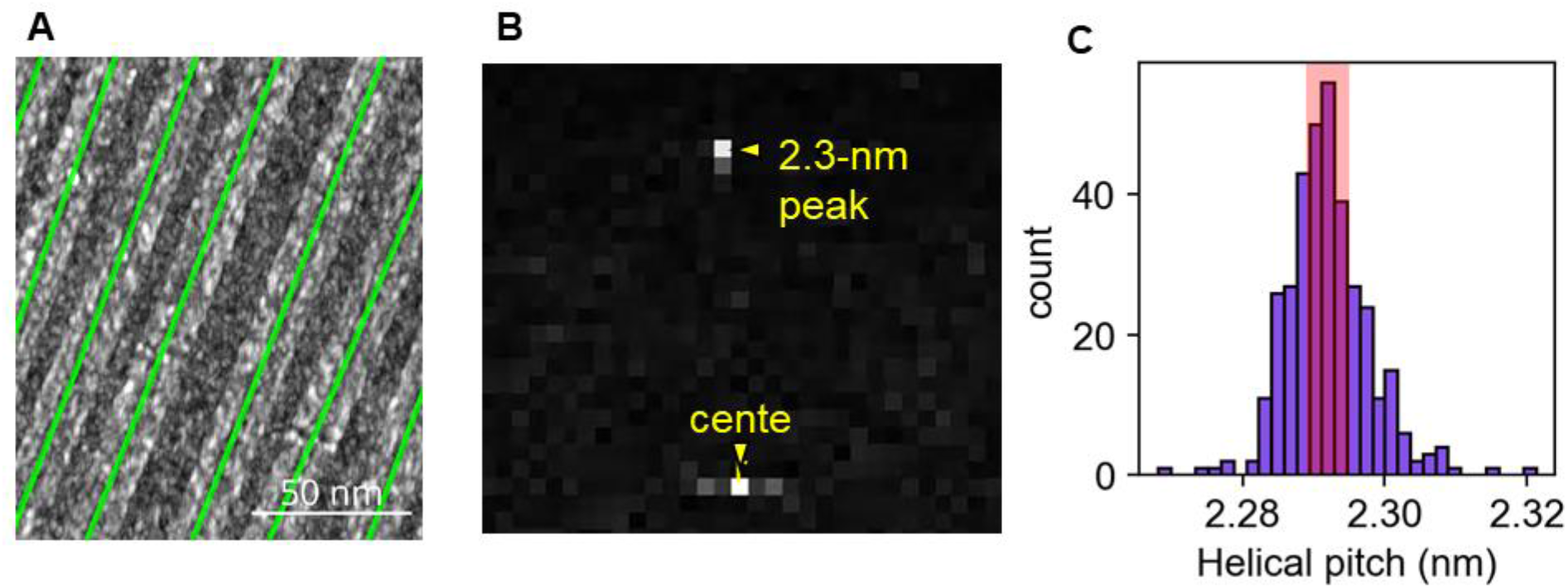
Pixel size calibration using local-CFT and tobacco mosaic viruses (TMVs). (A) A representative tomogram of TMVs and the fitted spline curves. (B) A representative cylindric power spectrum of a 46-nm TMV fragment. (C) A histogram of the helical pitch of each 46-nm TMV fragment normalized to the helical pitch value 2.292 ± 0.003 nm (shaded range) measured by X-ray fiber diffraction (Kendall et al., 2007).

**Fig. S2:**
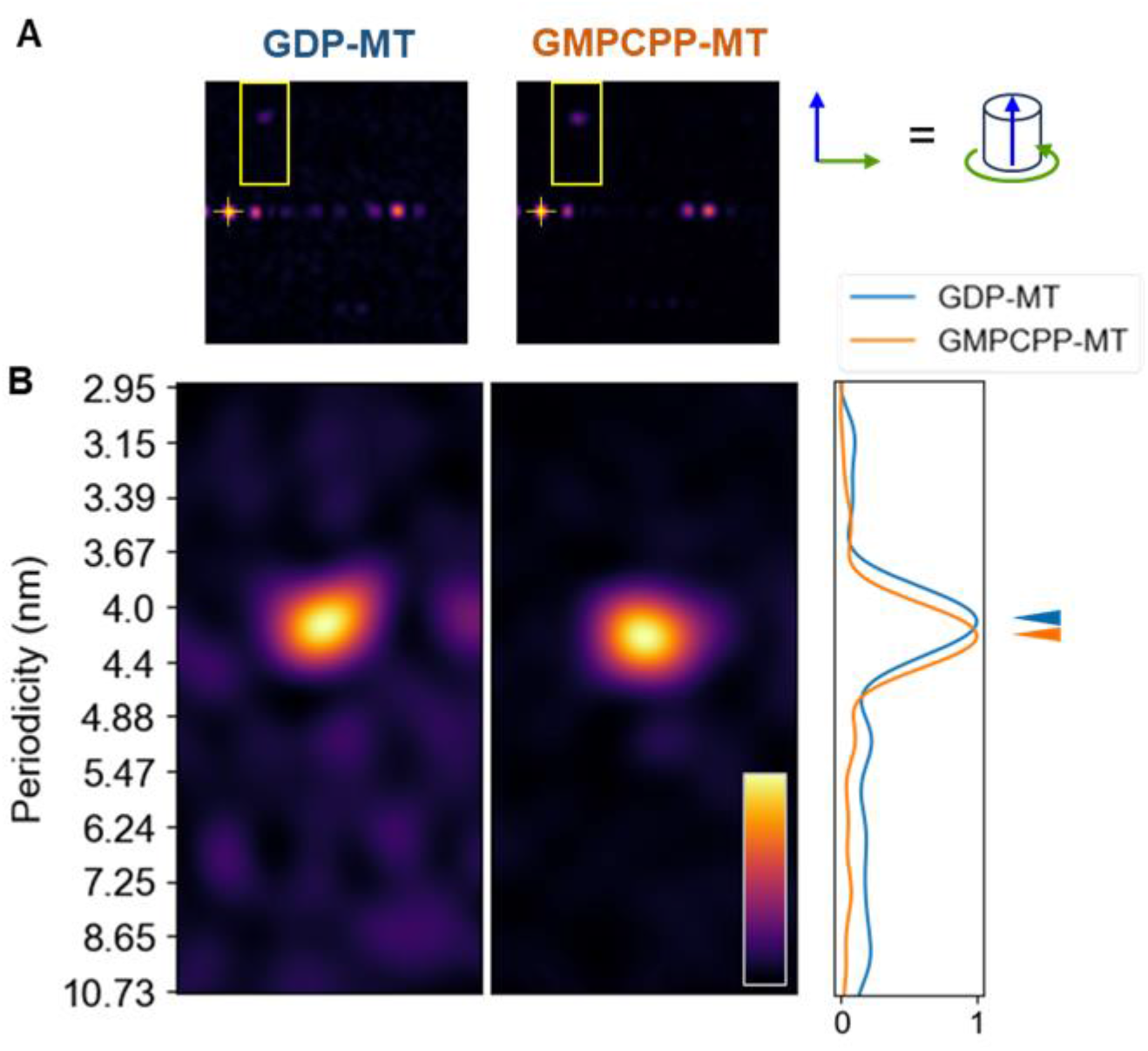
Examples of up-sampled cylindric power spectra of GDP-MT and GMPCPP-MT. (A) Power spectra around the center (shown as cross symbols). The y-axis corresponds to the longitudinal axis, and the x-axis corresponds to the angular axis. (B) Zoomed power spectra around the peaks that were used to estimate the longitudinal helical pitch parameters. The mean projection of each peak is shown on the right. A peak difference between GDP- and GMPCPP-MT is clearly detectable.

**Fig. S3:**
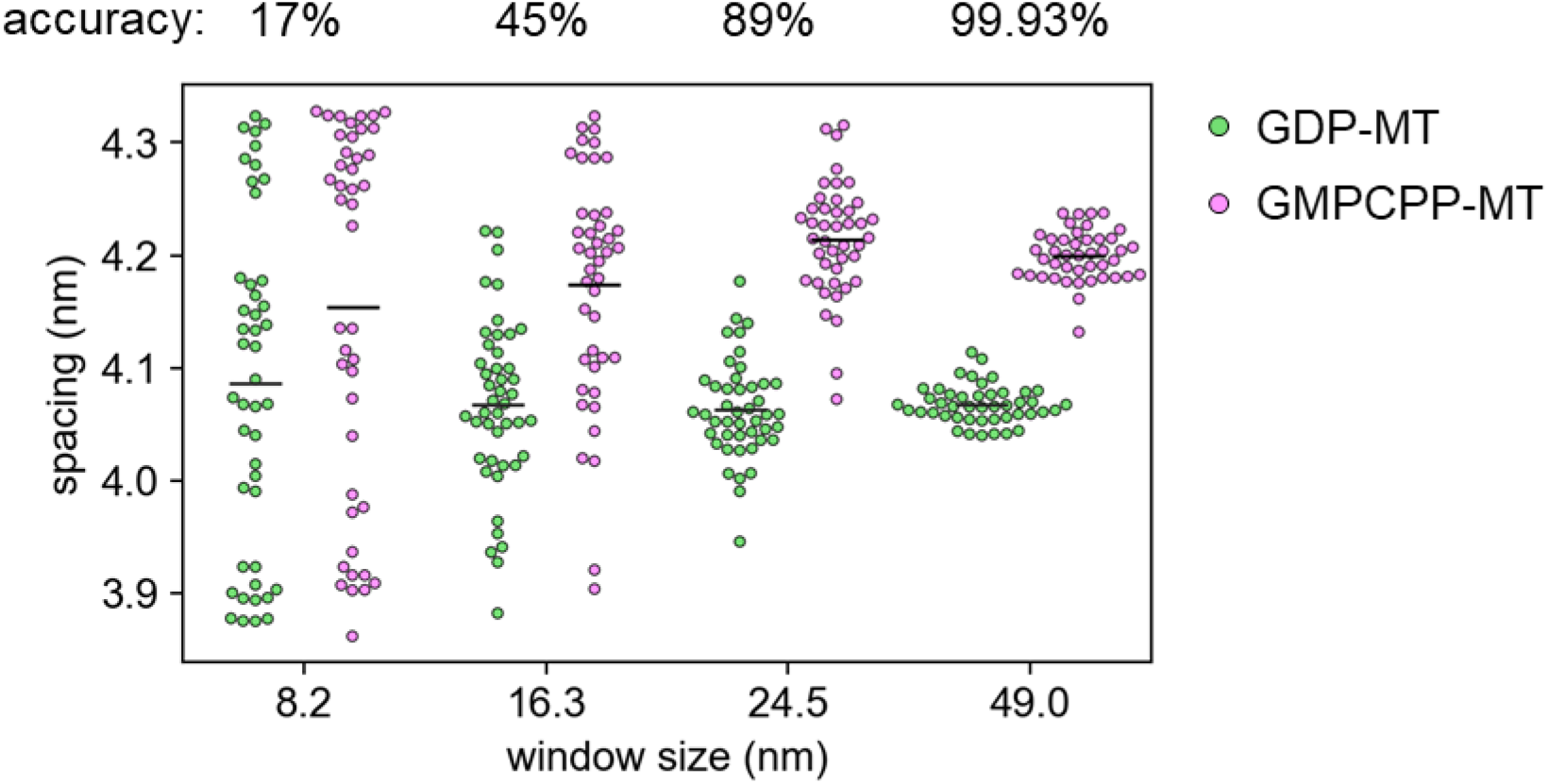
Relationship of CFT window sizes and accuracy of the microtubule type detection. Horizontal bars indicate mean values. Three microtubules were analyzed for each type: 45 fragments for GDP-MTs and 43 fragments for GMPCPP-MTs. The accuracy was calculated by the overlapping area of the two distributions, assuming the estimation error follows a normal distribution.

**Fig. S4:**
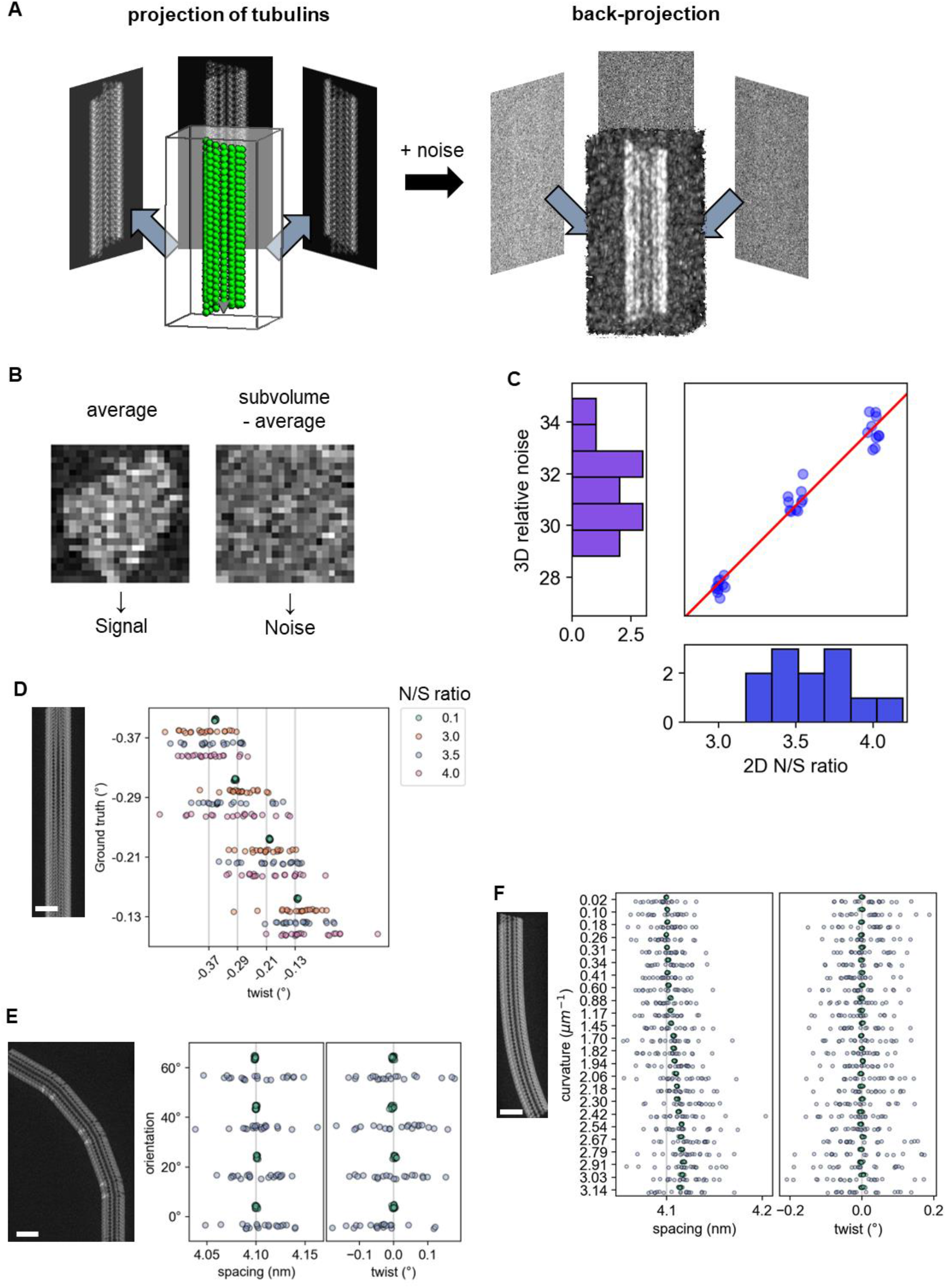
Tomogram simulation and noise calibration. (A) Schematic illustration of the tomogram simulation. (B) Estimation of the N/S ratio in tomograms using subtomogram averaging. (C) Calibration of the N/S ratio in the tilt series and the reconstructed tomogram. (D-F) A CFT analysis of simulated microtubules. Scale bars, 25 nm. (D) A 14_3 microtubule with twist angles of around 0.25°. (E) A 13_3 microtubule tilted 0 – 60° from the tilt axis (y-axis). The spacing (4.10 nm) and twist angle (0.0°) are constant. (F) A 13_3 microtubule with variable curvature. The curvature was calculated from the spline used for simulating the microtubule. The spacing (4.10 nm) and twist angle (0.0°) were constant.

**Fig. S5:**
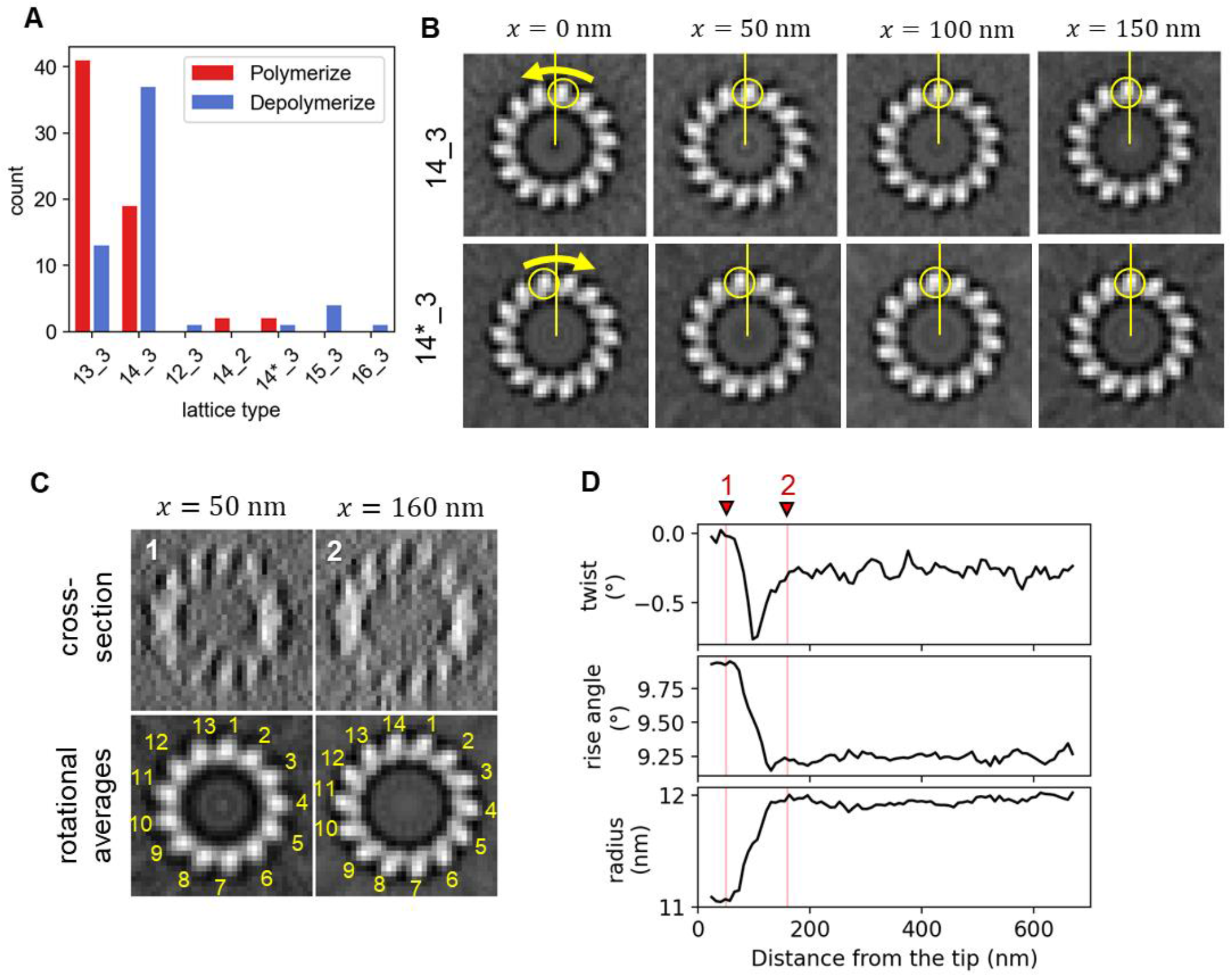
Microtubule lattice types. (A) Histogram of the distribution of microtubule N_S structures. (B) Comparison of ordinary 14_3 and rare 14*_3 microtubules. Images are rotationally averaged in 2D. (C) A microtubule from the depolymerizing sample with a change point of the protofilament number. (D) Profile of the local twist, rise angle and radius along the microtubule in C.

**Fig. S6:**
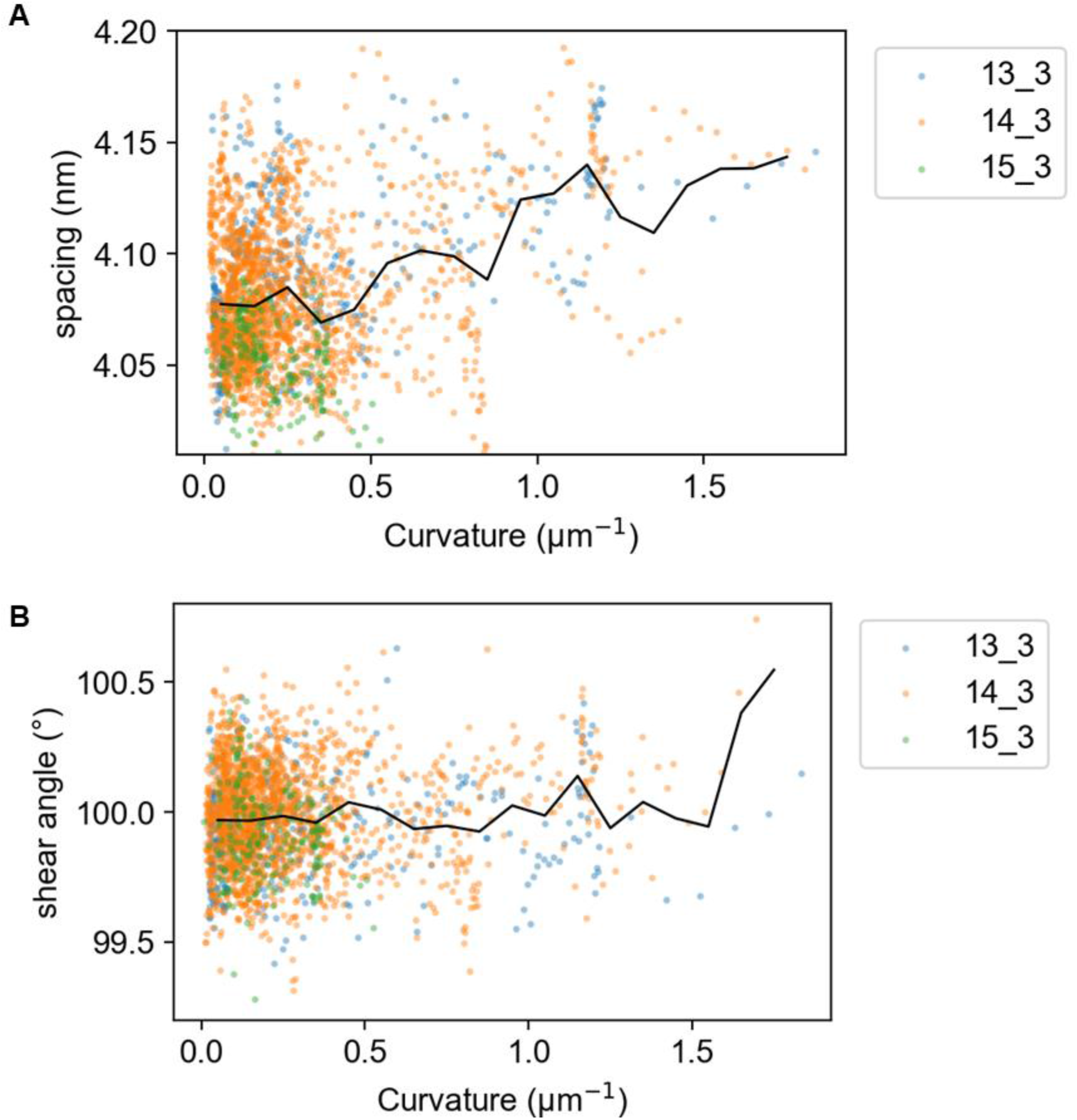
Dependency of lattice spacing (A) and shear angle (B) on the local curvature measured by CFT. The black line shows piecewise averages for each 0.1 μm^-1^ window. Some outliers are out of the figure region to highlight the change of the average lines.

**Fig. S7:**
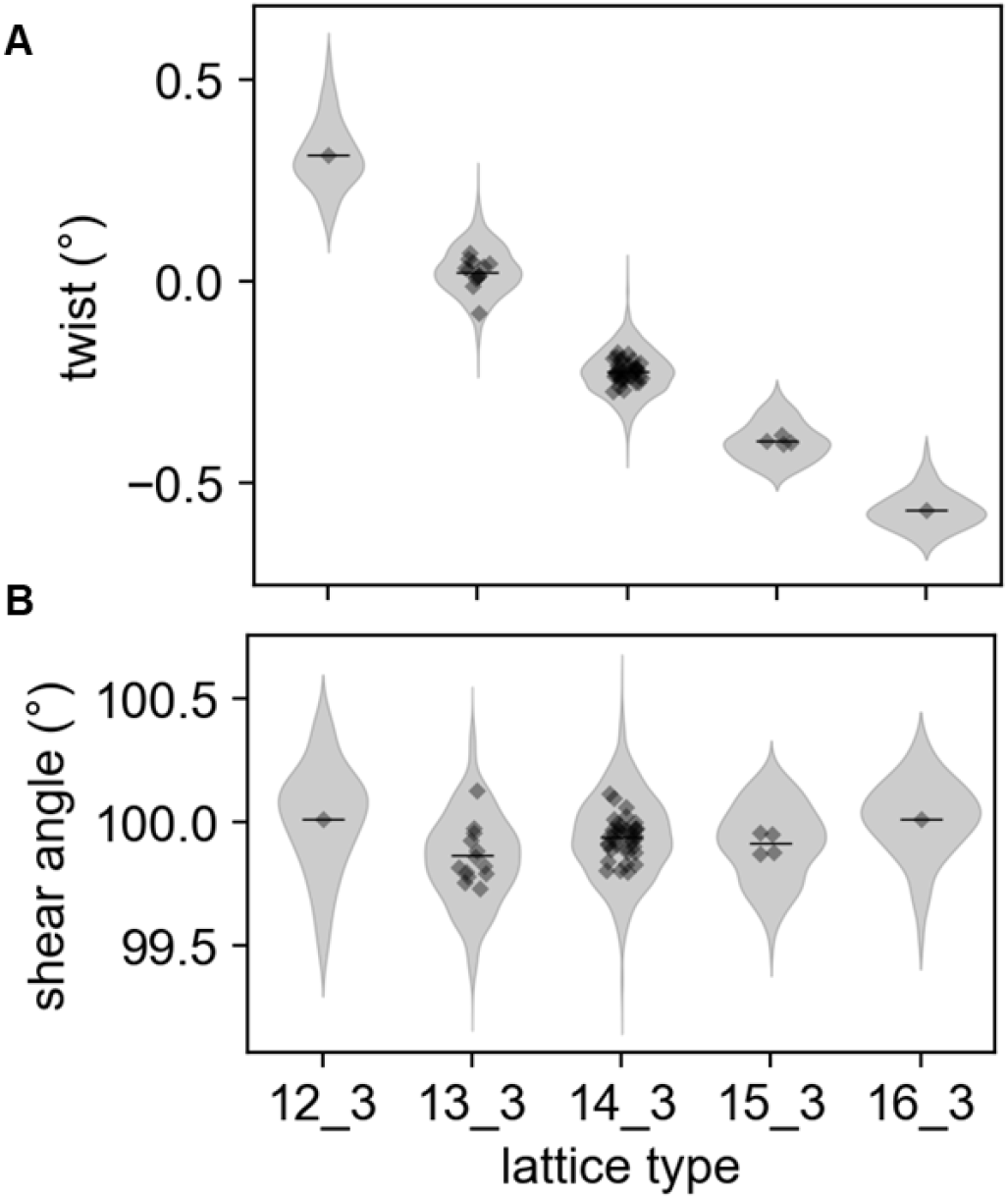
Distribution of (A) twist angles and (B) shear angles of depolymerizing microtubules of different lattice types.

**Fig. S8:**
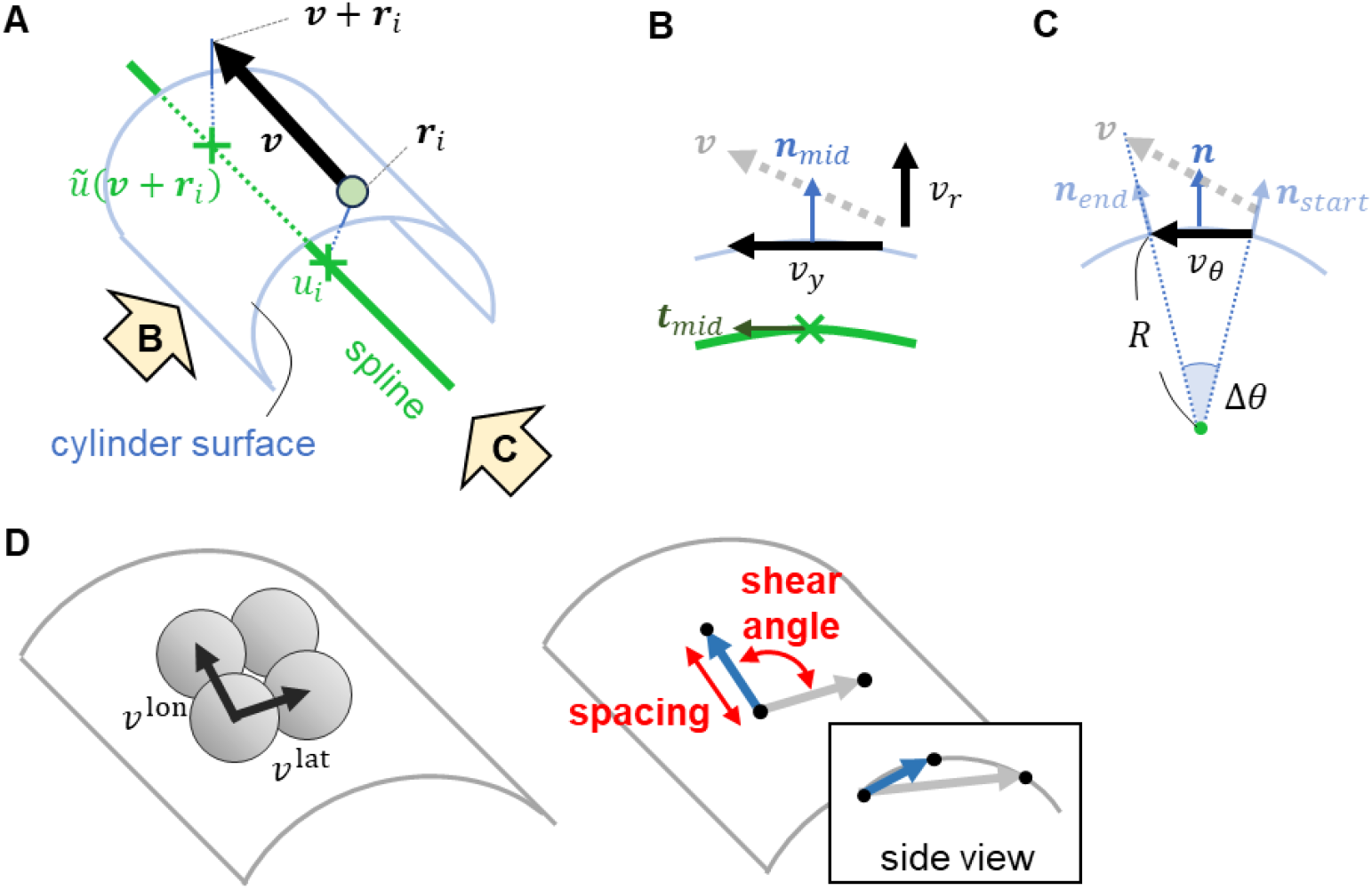
Method to calculate lattice parameters using the alignment results. (A) A cylinder surface surrounding a spline at constant radius *R* can be defined. Any vector ***v*** near the cylinder surface can be converted into the cylindric coordinate system as (*v*_*r*_, *v*_*y*_, *v*_*θ*_). (B) Sideview of the cylinder. *v*_*r*_ and *v*_*y*_ are calculated by the projection to the normal surface and the spline tangent vector. (C) Another sideview. *v*_*θ*_ is calculated by nonlinearly projecting vector ***v*** to the surface. (D) Inter-molecule vectors are used to calculate lattice parameters such as spacing and shear angle.

**Fig. S9:**
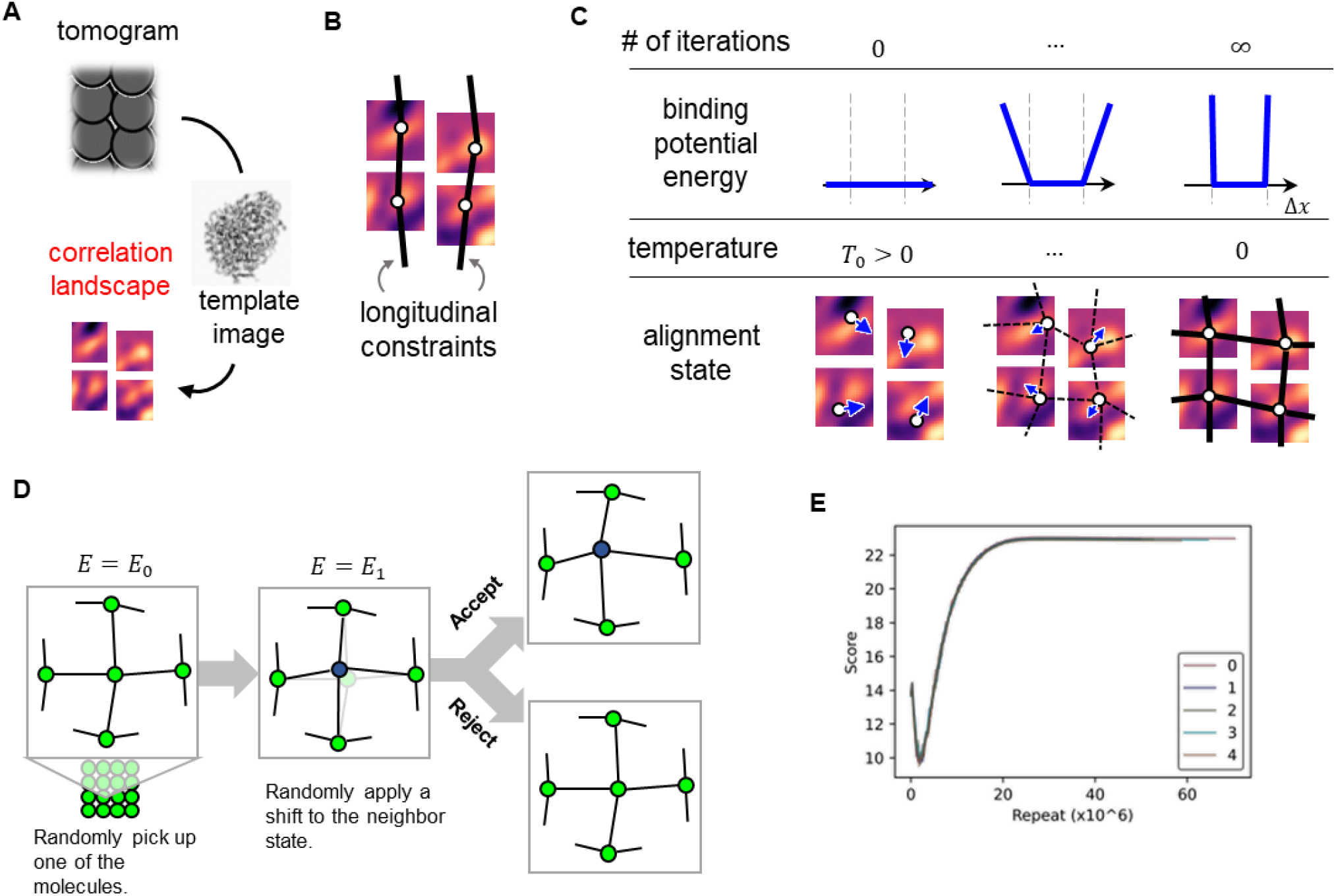
Schematic illustration of the constrained alignment algorithms. (A) First, the correlation landscape is built from the tomogram using tubulin template images. (B) In Viterbi alignment, only the longitudinal constraints are considered. Therefore, the total alignment score is optimized for each protofilament. (C) The RMA workflow. At the beginning, the binding potential is flat; thus, the inter-molecular penalty does not affect the score. The temperature is high enough to let all molecules freely transit to the neighbor state (blue arrows in the bottom image). As the iteration proceeds, the binding potential energy gets steeper, which means that the distances between tubulin neighbors are considered more rigid. At the same time, the temperature is cooled down, resulting in unfavorable state transitions being less frequent. (D) The detailed process of each iteration. One of the molecules is randomly shifted to one of the nearest positions, and the change in the “energy” is calculated. Whether to accept the change is determined depending on the temperature. (E) A representative convergence of the alignment score. Since RMA is a random process, five alignment tasks are conducted in parallel, and the best run is chosen as the solution.

**Fig. S10:**
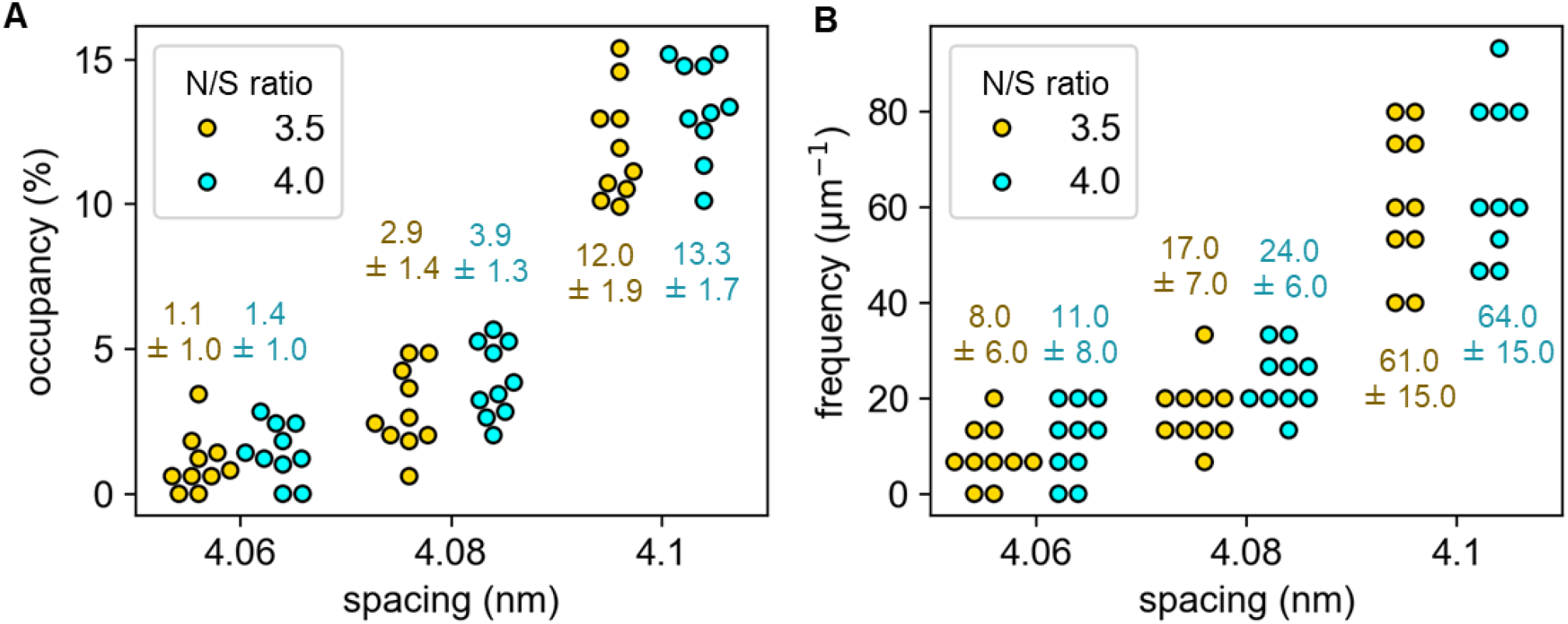
Lattice spacing heterogeneity analysis of simulated microtubules with constant lattice spacing. Three types of spacings (4.08 nm, 4.10 nm or 4.12 nm) and two N/S ratios (3.5 and 4.0) were used.

**Fig. S11:**
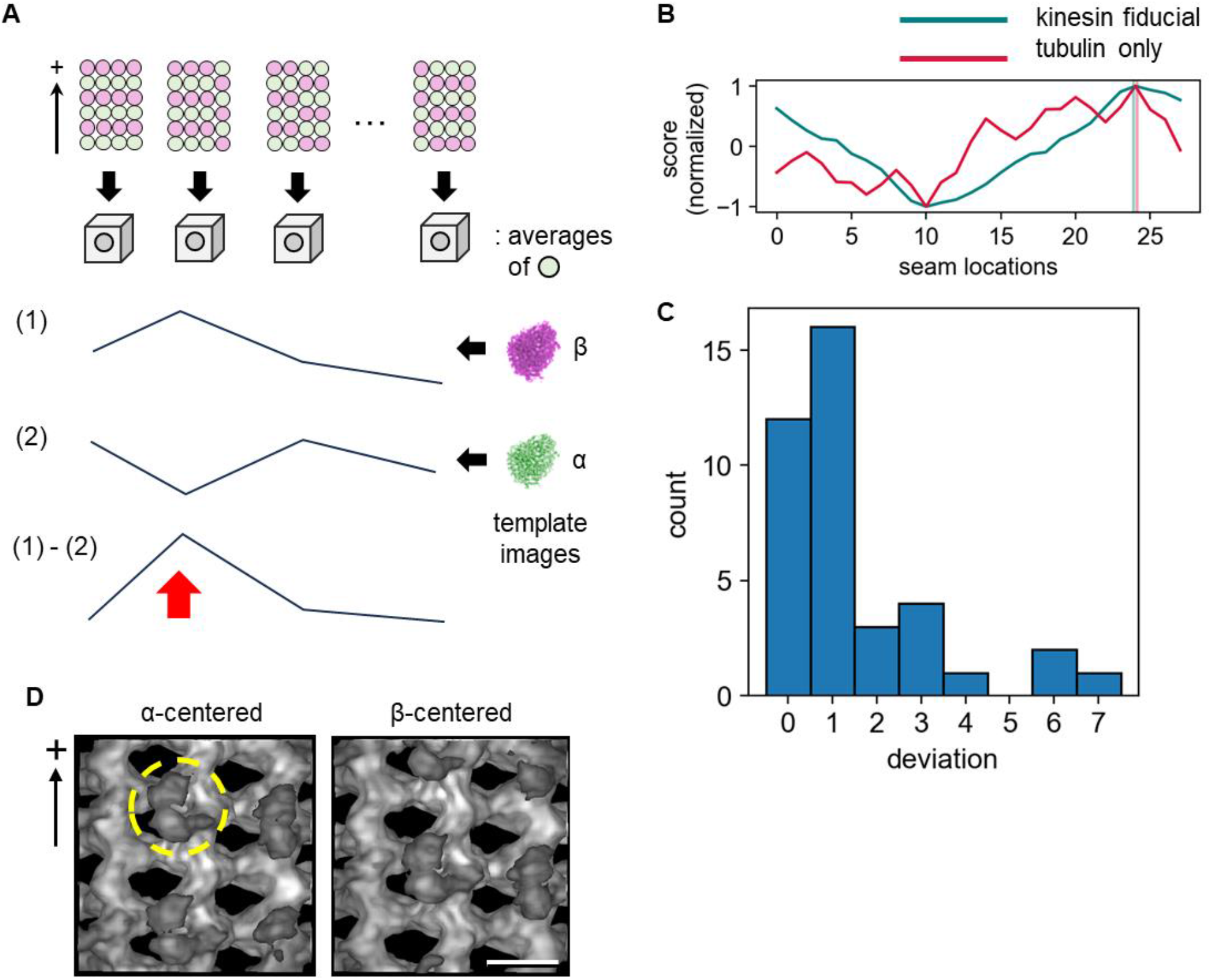
Seam-searching on kinesin-bound GMPCPP-MTs. (A) Schematic illustration of the seam-search protocol. For a microtubule with *N* protofilaments, there are 2*N* patterns of α/β-tubulin labeling. The average image is calculated for each pattern, and its correlation with an α- or β-tubulin template image is calculated. By subtracting these two correlations, the seam search score is obtained. The pattern that maximizes this score is used as the estimation of the seam location. (B) An example of the seam-search. To precisely estimate the seam location, kinesin-1 molecules bound to the GMPCPP-MT were used as the fiducials, and the kinesin-1 template image was used for the score calculation instead of the tubulin template image. (C) Distribution of the deviation between the correct seam location estimated using kinesin-1 and the location estimated by the fiducial-free seam-search. (D) Gaussian-filtered subtomogram averages around the estimated α- and β-tubulins. 6407 molecules were used in total. The yellow dashed circle indicates a kinesin-1 molecule. Scale bar, 5 nm.

**Fig. S12:**
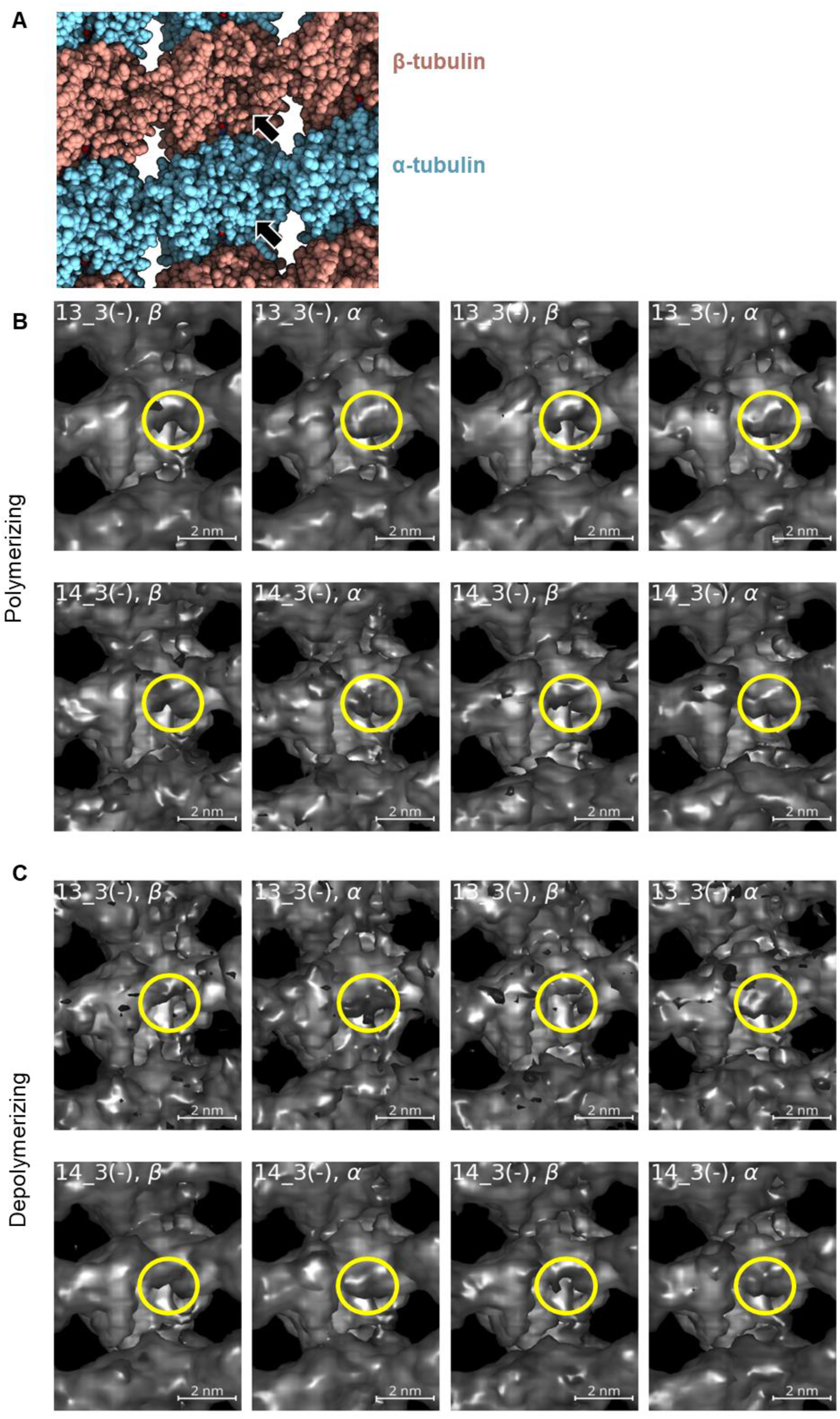
Results of the seam-search. (A) An atomic model of αβ-tubulins in a microtubule (PDB: 6DPV; rendered by ChimeraX (Goddard et al., 2018)). Arrows indicate the distinctive difference between α- and β-tubulin; that is, the loop region only seen with α-tubulins. (B, C) Subtomogram averages of polymerizing/depolymerizing, plus-end/minus-end, and 13_3/14_3 microtubules from the luminal side. An α- or β-tubulin was placed in the center of each panel. Yellow circles correspond to the density shown in (A). For each panel, the top side is oriented at the minus end. The same threshold value and contrast were used to render all images.

### Tables

**Table S1:** Lattice parameters of different types of microtubules measured by local-CFT (mean ± S. D.).

| Microtubule type | N_S | Spacing (nm) | Twist ($^{\circ}$) | Rise length (nm) | # of fragments |
| --- | --- | --- | --- | --- | --- |
| GDP-MT | 13_3 | $4.079 \pm 0.026$ | $0.031 \pm 0.066$ | $0.942 \pm 0.006$ | 97 |
| | 14_3 | $4.076 \pm 0.024$ | $-0.204 \pm 0.054$ | $0.871 \pm 0.005$ | 160 |
| | 15_3 | $4.070 \pm 0.021$ | $-0.355 \pm 0.089$ | $0.817 \pm 0.028$ | 31 |
| GMPCPP-MT | 13_3 | $4.216 \pm 0.021$ | $0.078 \pm 0.110$ | $0.971 \pm 0.038$ | 48 |
| | 14_3 | $4.200 \pm 0.027$ | $-0.173 \pm 0.074$ | $0.899 \pm 0.014$ | 671 |
| | 16_4 | $4.205 \pm 0.017$ | $0.418 \pm 0.050$ | $1.061 \pm 0.005$ | 17 |

**Table S2:** Estimation of lattice spacing of simulated microtubules by local-CFT (mean ± S. D.).

| Ground truth (nm) | Estimated lattice spacing (nm) |  |  |  |
| --- | --- | --- | --- | --- |
|  | N/S = 0.1 | N/S = 3.0 | N/S = 3.5 | N/S = 4.0 |
| 4.05 | $4.0504 \pm 0.0005$ | $4.055 \pm 0.023$ | $4.047 \pm 0.030$ | $4.044 \pm 0.032$ |
| 4.10 | $4.1010 \pm 0.0005$ | $4.103 \pm 0.017$ | $4.103 \pm 0.020$ | $4.103 \pm 0.038$ |
| 4.15 | $4.1510 \pm 0.0007$ | $4.153 \pm 0.014$ | $4.149 \pm 0.032$ | $4.147 \pm 0.030$ |
| 4.20 | $4.1999 \pm 0.0006$ | $4.198 \pm 0.026$ | $4.195 \pm 0.024$ | $4.189 \pm 0.032$ |

**Table S3:** Estimation of twist angles of simulated 13_3 microtubules by local-CFT (mean ± S. D.).

| Ground truth ($^{\circ}$) | Estimated twist angle ($^{\circ}$) | | | |
| --- | --- | --- | --- | --- |
|  | N/S = 0.1 | N/S = 3.0 | N/S = 3.5 | N/S = 4.0 |
| -0.12 | $-0.115 \pm 0.002$ | $-0.112 \pm 0.065$ | $-0.120 \pm 0.071$ | $-0.129 \pm 0.067$ |
| -0.04 | $-0.039 \pm 0.002$ | $-0.062 \pm 0.066$ | $-0.014 \pm 0.056$ | $-0.003 \pm 0.064$ |
| 0.04 | $0.040 \pm 0.002$ | $0.000 \pm 0.062$ | $0.037 \pm 0.068$ | $0.012 \pm 0.098$ |
| 0.12 | $0.116 \pm 0.002$ | $0.126 \pm 0.055$ | $0.114 \pm 0.071$ | $0.098 \pm 0.108$ |

**Table S4:**
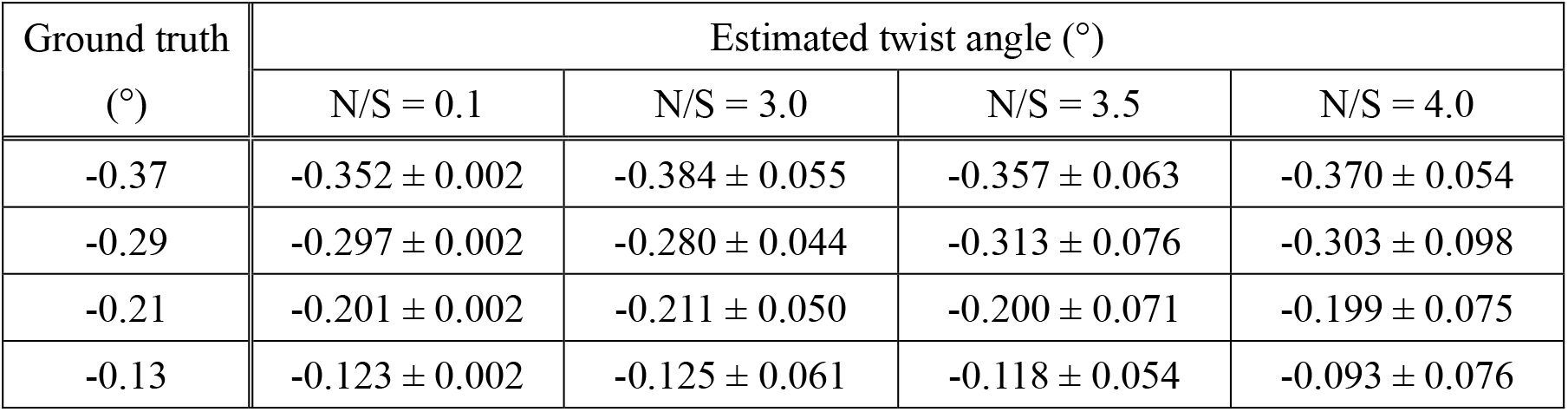
Estimation of twist angles of simulated 14_3 microtubules by local-CFT.

| Ground truth ($^{\circ}$) | Estimated twist angle ($^{\circ}$) | | | |
| --- | --- | --- | --- | --- |
|  | N/S = 0.1 | N/S = 3.0 | N/S = 3.5 | N/S = 4.0 |
| -0.37 | $-0.352 \pm 0.002$ | $-0.384 \pm 0.055$ | $-0.357 \pm 0.063$ | $-0.370 \pm 0.054$ |
| -0.29 | $-0.297 \pm 0.002$ | $-0.280 \pm 0.044$ | $-0.313 \pm 0.076$ | $-0.303 \pm 0.098$ |
| -0.21 | $-0.201 \pm 0.002$ | $-0.211 \pm 0.050$ | $-0.200 \pm 0.071$ | $-0.199 \pm 0.075$ |
| -0.13 | $-0.123 \pm 0.002$ | $-0.125 \pm 0.061$ | $-0.118 \pm 0.054$ | $-0.093 \pm 0.076$ |

**Table S5:** Lattice parameters of polymerizing and depolymerizing microtubules measured by local-CFT (mean ± S. D.).

| Lattice type | Tip type and state |  | Spacing (nm) | Shear angle (°) |
| --- | --- | --- | --- | --- |
| 13_3 | Plus | Polymerizing | $4.079 \pm 0.025$ | $100.00 \pm 0.19$ |
| | | Depolymerizing | $4.058 \pm 0.019$ | $99.84 \pm 0.18$ |
| | Minus | Polymerizing | $4.069 \pm 0.021$ | $99.89 \pm 0.18$ |
| | | Depolymerizing | $4.058 \pm 0.015$ | $99.84 \pm 0.18$ |
| | Middle region | | $4.082 \pm 0.027$ | $99.91 \pm 0.19$ |
| 14_3 | Plus | Polymerizing | $4.080 \pm 0.017$ | $100.05 \pm 0.17$ |
| | | Depolymerizing | $4.066 \pm 0.024$ | $99.92 \pm 0.19$ |
| | Minus | Polymerizing | $4.078 \pm 0.021$ | $99.96 \pm 0.14$ |
| | | Depolymerizing | $4.067 \pm 0.021$ | $99.93 \pm 0.17$ |
| | Middle region | | $4.080 \pm 0.030$ | $99.99 \pm 0.18$ |

**Table S6:** Linear regression coefficients of curvature dependency of microtubule lattice parameters.

| Type | Region | Slope |  |
| --- | --- | --- | --- |
| | | Spacing (nm/$\mu\text{m}^{-1}$) | Twist ($^{\circ}/\mu\text{m}^{-1}$) |
| 13_3 | Inner | $-0.006 \pm 0.009$ | $-0.000 \pm 0.043$ |
| | Side | $0.026 \pm 0.010$ | $-0.002 \pm 0.042$ |
| | Outer | $0.056 \pm 0.009$ | $0.006 \pm 0.045$ |
| 14_3 | Inner | $-0.005 \pm 0.009$ | $0.003 \pm 0.037$ |
| | Side | $0.028 \pm 0.009$ | $-0.001 \pm 0.033$ |
| | Outer | $0.063 \pm 0.009$ | $0.006 \pm 0.034$ |

